# Resistance, heteroresistance and fitness costs drive colistin treatment failure during *Acinetobacter baumannii* pneumonia

**DOI:** 10.1101/2025.07.24.666669

**Authors:** Juan Hernandez-Bird, Bixi He, Leah M. VanOtterloo, Elizabeth B. Billings, Wenwen Huo, Gabriella I C Teodoro, Haley Echlin, Jason M. Rosch, M. Stephen Trent, Ralph R. Isberg

**Affiliations:** Department of Molecular Biology and Microbiology, Tufts University School of Medicine, MA, USA; Department of Microbiology, College of Arts and Sciences, University of Georgia, GA, USA; Department of Infectious Diseases, St Jude Children’s Research Hospital, TN, USA

## Abstract

*Acinetobacter baumannii* is an ESKAPE pathogen linked to healthcare-associated diseases. Due to evolved resistance, last-resort antibiotics such as the lipooligosaccharide (LOS)-targeting colistin are increasingly used to treat multidrug-resistant isolates. To track the evolution of colistin resistance within a host, we performed sequential oropharyngeal infections in immunocompetent or immune-depleted mice in the presence of inhaled colistin. Both resistant and heteroresistant *A. baumannii* strains emerged with *pmrB* mutations that efficiently competed with the susceptible parent in the presence of colistin. These *pmrB* mutants had a fitness cost in the absence of colistin treatment but retained their ability to colonize the host. In contrast, LOS-deficient *A. baumannii* mutants removed the target of colistin, but such mutants were unable to colonize the lung. The two pathogenic *pmrB* mutants showed clear evidence of LOS modification, which was linked to increased transcription of LOS modification enzymes, including the product of the cryptic *eptA* gene. Spontaneous insertion mutations that caused hyperexpression of *eptA* allowed the heteroresistant mutant to develop clinically-significant colistin resistance. Insertion mutations upstream of the *eptA* gene or those disrupting *hns*, which encodes a small histone-like protein, resulted in increased *eptA* transcript, linking expression of this protein to clinically significant resistance. A resistant variant derived from the heteroresistant parent was stable in the absence of drug, but continued passaging selected for colistin-resensitized pseudorevertants that were largely due to disruption of the LOS modification enzymes. Therefore, colistin heteroresistance is an early stage in the stepwise acquisition of stable colistin resistance in *A. baumannii*.

**Significance:** The mutational pathways leading to antibiotic-resistant infections and the role of the immune system in preventing them are poorly understood. Here we employed a colistin-treated mouse pneumonia model of *Acinetobacter baumannii* and observed the evolution of colistin-resistant and heteroresistant mutants in immune-depleted and immunocompetent hosts, respectively. We show that mutations that result in alteration of colistin’s target drive evolutionary pathways to resistance, whereas removal of the target is unlikely to be clinically significant. We demonstrated that the heteroresistant mutant generates subpopulations with higher levels of resistance through insertion mutations in specific gene regions. This study furthers our understanding of how resistance emerges during infection and provides a genetic explanation for the transition from colistin heteroresistance to full resistance.

## Introduction

Antibiotic-resistant infections caused by the ESKAPE group of pathogens are an urgent global health concern, with lethality predicted to surpass cancer by 2050(1). Belonging to this group is *Acinetobacter baumannii*, a Gram-negative bacterium with a remarkable ability to evolve resistance to currently employed antibiotics(2). As an opportunistic pathogen, *A. baumannii* mainly affects wounded or immunocompromised patients in healthcare settings with respiratory, bloodstream and urinary tract infections being its most concerning clinical manifestations(3, 4). The World Health Organization currently classifies carbapenem-resistant *A. baumannii* in the highest priority category for research and development of treatment options(5). Therefore, understanding how this organism evolves to give rise to drug-resistant variants is of the upmost importance.

Resistance to the first-line antimicrobial carbapenems is increasingly common in *A. baumannii* clinical isolates(6), so last-resort antibiotics are currently being employed against extensively drug-resistant *A. baumannii*(7). Colistin is one such antimicrobial, but its usefulness is limited due to concerns about its nephrotoxicity. New inhalation therapies are being pursued for the treatment of respiratory infections in hopes of reducing toxic side effects(8, 9). The drug is a positively charged peptide that binds to the negatively charged phosphate groups on the lipid A component of lipopolysaccharide (LPS), or lipooligosaccharide (LOS) in the case of *A. baumannii,* leading to envelope disruption and cell lysis(7).

Gram-negative bacteria have evolved multiple strategies to modify the lipid A component of LPS/LOS to increase resistance to antimicrobial peptides(10). In *A. baumannii,* colistin resistance is mainly caused by mutations within the *pmrCAB* operon(11). The PmrAB cassette is a two-component system that responds to low pH and select divalent cations (Fe^2+^, Zn^2+^, Al^3+^), controlling the expression of multiple genes including the *pmrC* and the *naxD* operons(12). *pmrC,* also known as *eptA*, encodes a phosphoethanolamine transferase that adds phosphoethanolamine to the 1- and 4’ phosphoryl groups in lipid A(13). *naxD* encodes a YdjC family deacetylase that plays a part in the addition of galactosamine to the 1-phosphoryl group in lipid A(14). These modifications reduce both the negative charge and LOS affinity for polymyxins such as colistin(13). Mutations in the *pmrAB* genes can lead to the constitutive activation of this system by the PmrA positive regulator, resulting in polymyxin resistance. In addition, some strains of *Acinetobacter* possess an orphan homologue of the PmrC/EptA phosphoethanolamine transferase, that is also under the control of PmrAB. For simplicity, we will refer to the orphan copy as *eptA* and the *pmrAB-*linked copy as *pmrC.* Clinical isolates with high colistin resistance have been characterized that overexpress *eptA* transcript as a consequence of insertion sequence transposition into regions upstream of *eptA*(15, 16).

Mutations in *pmrAB* have also been linked to colistin heteroresistance in *A. baumannii*(17). Antibiotic heteroresistant strains generate subpopulations resistant to the exposed antibiotic at much higher frequencies than observed for sensitive strains(18). Heteroresistant pathogens are troubling because current clinical microbiology methods frequently classify them as susceptible, and their genetic background could provide an intermediate step towards the evolution of traditional resistance(19–21). The genetic mechanisms facilitating this phenomenon are poorly understood, but recent evidence points to transient gene amplification in the resistant subpopulation, which occurs at a higher frequency than observed for spontaneous mutations(22). Amplification and mutations in genes that regulate or mediate the modification of lipid A have been reported to contribute to colistin heteroresistance in pathogens other than *Acinetobacter*(23–26). The genetic basis for colistin heteroresistance in *A. baumannii*, however, appears unrelated to amplification.

*A. baumannii* is one of the few known species with the ability to survive with an LOS-deficient outer membrane(27). LOS-deficient mutants are readily isolated in a subset of clinical isolates during *in-vitro* experimental evolution to polymyxins but are rarely reported in clinical studies(28). Absence of LOS, usually due to mutations in the early LOS biosynthesis genes, results in high-level colistin resistance but is accompanied by outer membrane instability and a high fitness cost(29). Nevertheless, these LOS-deficient mutants may provide transition states to more virulent, drug-resistant mutants(30, 31). Inactivation of PBP1A glycosyltransferase activity allows viability in the presence of LOS knockout mutations(32). Additionally, mutations in the *pldA* and *mla* genes, enzymes that act to reduce glycerophospholipid content in the outer membrane, alleviate the severe growth defect in LOS-deficient strains(27). It is unknown if these suppressor mutations allow LOS-deficient strains to retain their virulence.

Previously, we performed serial passaging of the low virulence *A. baumannii* ATCC17978 strain in a murine pneumonia model during treatment with ciprofloxacin(33). In immune-depleted animals, drug-tolerant/persistent mutants arose that resulted in antibiotic treatment failure and the emergence of second-step drug-resistant mutants. Neither class of mutants achieved high population levels in the lines passaged in immunocompetent hosts(33, 34). More recent multi-drug resistant *A. baumannii* clinical isolates have been proposed as more relevant for infection modeling due to their ability to cause lethal pneumonia even in the absence of immune depletion(35). One of these strains is *A. baumannii* LAC-4, a multi-drug resistant specimen that was isolated from a Los Angeles County outbreak in 1997(36–38),. In this study, we performed experimental evolution in this highly mouse-virulent strain, identifying multiple routes to colistin-resistant virulent isolates, as well as forbidden routes that result in avirulent strains.

## Results

### Sequential lung infections in the presence of nebulized colistin select for pmrB mutations

To better simulate the conditions in which *A. baumannii* evolves colistin resistance, we performed 16 sequential lung infections with the LAC-4 strain in both immune-depleted and immunocompetent mice treated with inhaled colistin (Fig. 1A). Immune-depleted animals received two intraperitoneal injections of cyclophosphamide, a chemotherapeutic agent that induces profound neutropenia and significantly reduces lymphocyte and monocyte counts(39, 40). Lung infections were established via oropharyngeal inoculation in both sets of animals. Three separate lines were maintained per condition, and lung bacterial burden after 24-hour infections was quantified (Figs. 1B, C). The inoculum CFU for the immune-depleted lines was reduced in the second and fifth passage to ensure effective antibiotic selection throughout the experiment (2 x 10^2^ CFU below the expected bacterial burden; Fig. 1B). Line A showed evidence of antibiotic treatment failure after 13 passages in the immune-depleted animals, with bacterial burdens approaching that observed in untreated animals. In contrast, colistin treatment consistently reduced bacterial burden in the other two immune-depleted lines and in the immunocompetent lines throughout all passages (Figs. 1B, C).

**Figure 1:**
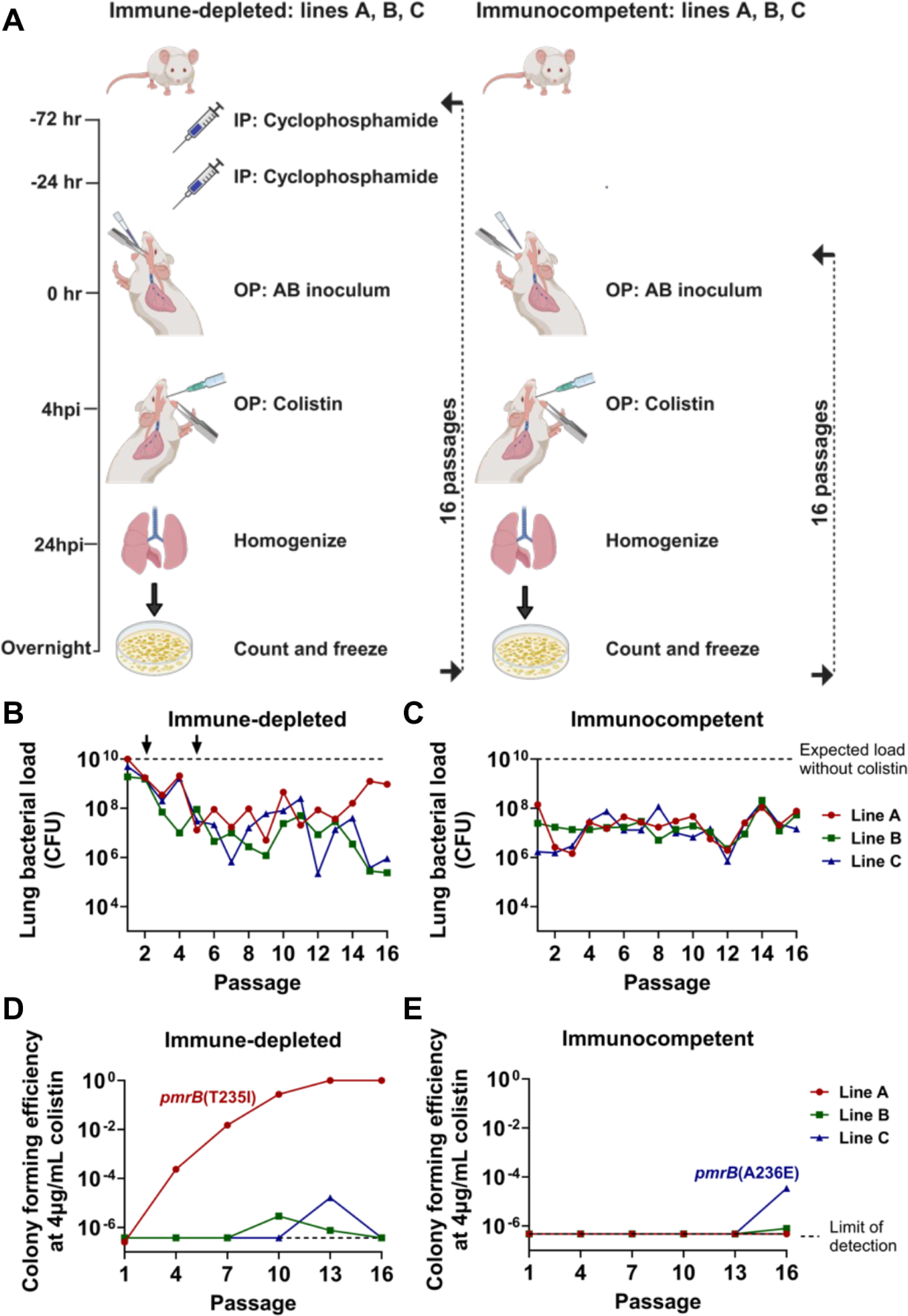
Colistin treatment selects for *pmrB* mutations in *A. baumannii* during serial oropharyngeal inoculations. **A.** Passaging strategy. Immune depletion was induced by intraperitoneal (IP) cyclophosphamide injections at 72- and 24-hours before infection. Three separate, co-housed lines derived from different LAC-4 colonies were passaged per condition. *A. baumannii* LAC-4 was oropharyngeally (OP) inoculated into mice and 4 hours post-infection (hpi), 8mg/kg aerosolized colistin was administered via OP inhalation. At 24 hpi, the mice were euthanized, lungs were removed, homogenized, and plated on LB agar plates. After overnight growth, the bacteria were collected and saved for the next round of infections. **B-C.** Lung CFU after each passage in colistin-treated mice. Dotted line: expected bacterial burden in the absence of colistin treatment (Materials and Methods). At passage 2 in immune-depleted host, the inoculum size of 5×10^7^ CFU was reduced to 1×10^7^ CFU, followed by a reduction to 5×10^6^ CFU at passage 5 (arrows). Inoculum size in immunocompetent host: 1×10^8^ CFU. **D-E.** Identification of colistin-resistant mutants. Pools were serially diluted and plated on LB plates supplemented with 0 or 4 μg/mL colistin. Colony forming efficiency (CFE) was calculated by the ratio of colonies growing in the presence or absence of drug. Dotted line: limit of detection. Isolates that grew in colistin plates were whole-genome sequenced to reveal the presence of the noted *pmrB* mutations.

Colistin-resistant *A. baumannii* isolates are defined by the Clinical Laboratory Standards Institute (CLSI) as being able to grow at colistin concentrations ≥ 4μg/mL(41). The parental LAC-4 strain is colistin-susceptible, so saved pools were plated on LB agar supplemented with 0-4μg/mL colistin to detect the presence and abundance of colistin-resistant isolates (Figs. 1D, E and SI Appendix, Fig. S1). During passaging in the immune-depleted animals, colistin-resistant colonies emerged in all three lines, but only in line A did the proportion of resistant clones increase and overgrow the susceptible ancestor (Fig. 1D). Whole genome sequencing (WGS) of saved resistant isolates from line A revealed they all have the *pmrB(*T235I) mutation that has been previously described in polymyxin-resistant *A. baumannii* strains(42).

Colistin-resistant colonies emerged in lines B and C in the last three passages through immunocompetent animals (Fig. 1E). WGS of resistant isolates in line C revealed that they all possessed the *pmrB*(A236E) mutation that had been associated previously with polymyxin resistance(43). Due to their increase in proportion on solid medium containing 4μg/ml colistin on successive passages through mice, the *pmrB*(T235I) and *pmrB*(A236E) clones were selected for further characterization. Importantly, the *pmrB*(T235I) strain employed in this study contains three additional IS*Aba* insertion mutations discussed later on. Additional mutants appeared throughout the passaging that grew at low colistin concentrations. These were isolated, subjected to MIC determination and WGS, but they were not studied further (Dataset S1 and SI Appendix, Fig. S1). Thus, colistin treatment in both host immune states selected for *A. baumannii* isolates with mutations in the region encoding the PmrB histidine kinase domain during passage in the experimental pneumonia model.

### The fitness cost of the LAC-4 pmrB mutants

Mutations that promote antibiotic resistance are often associated with fitness costs(44, 45). The continuous spread of drug-resistant bacteria requires that resistant mutants compete efficiently with fully virulent parental organisms in mammalian tissues(46). Therefore, we tested the hypotheses that mutations selected during lung infections would confer resistance without compromising the ability of the LAC-4 to colonize the host, and that these mutants would overgrow the parental strain in the presence of colistin.

The two evolved *pmrB* mutants were compared to WT LAC-4 during growth in broth culture. The *pmrB*(T235I) mutant, which had been selected in the immune-depleted animal, had a significant increase in broth doubling time compared to WT (Fig. 2A; p≤0.05). In contrast, the *pmrB*(A236E) mutant that was selected in the immunocompetent animal showed no significant change in doubling time during broth growth (Fig. 2A). To determine the relative fitness of the evolved *pmrB* mutants compared to WT during lung infection, strains were mixed 1:1 (mutant:WT) and competition experiments were performed during experimental lung infections in the presence or absence of immune depletion (Figs. 2B, C). In the absence of colistin treatment, the evolved *pmrB* mutants exhibited a competition deficiency against WT in the immunocompetent host, with the *pmrB*(T235I) mutant exhibiting the larger defect (Fig. 2B). In contrast, there was no significant competition defect against WT when the competition was performed in immune-depleted hosts (Fig. 2C). The gentamycin-susceptible LAC-4 strain (*ΔpABLAC2*) was included in this study since it exhibits no significant change in colistin sensitivity and growth rate *in-vitro,* allowing us to perform WT vs WT mouse competitions as a control (Figs. 2A-C). These results demonstrate that in the absence of colistin, *A. baumannii pmrB* mutants experimentally evolved during lung infections can be negatively selected in the presence of an intact immune response.

**Figure 2:**
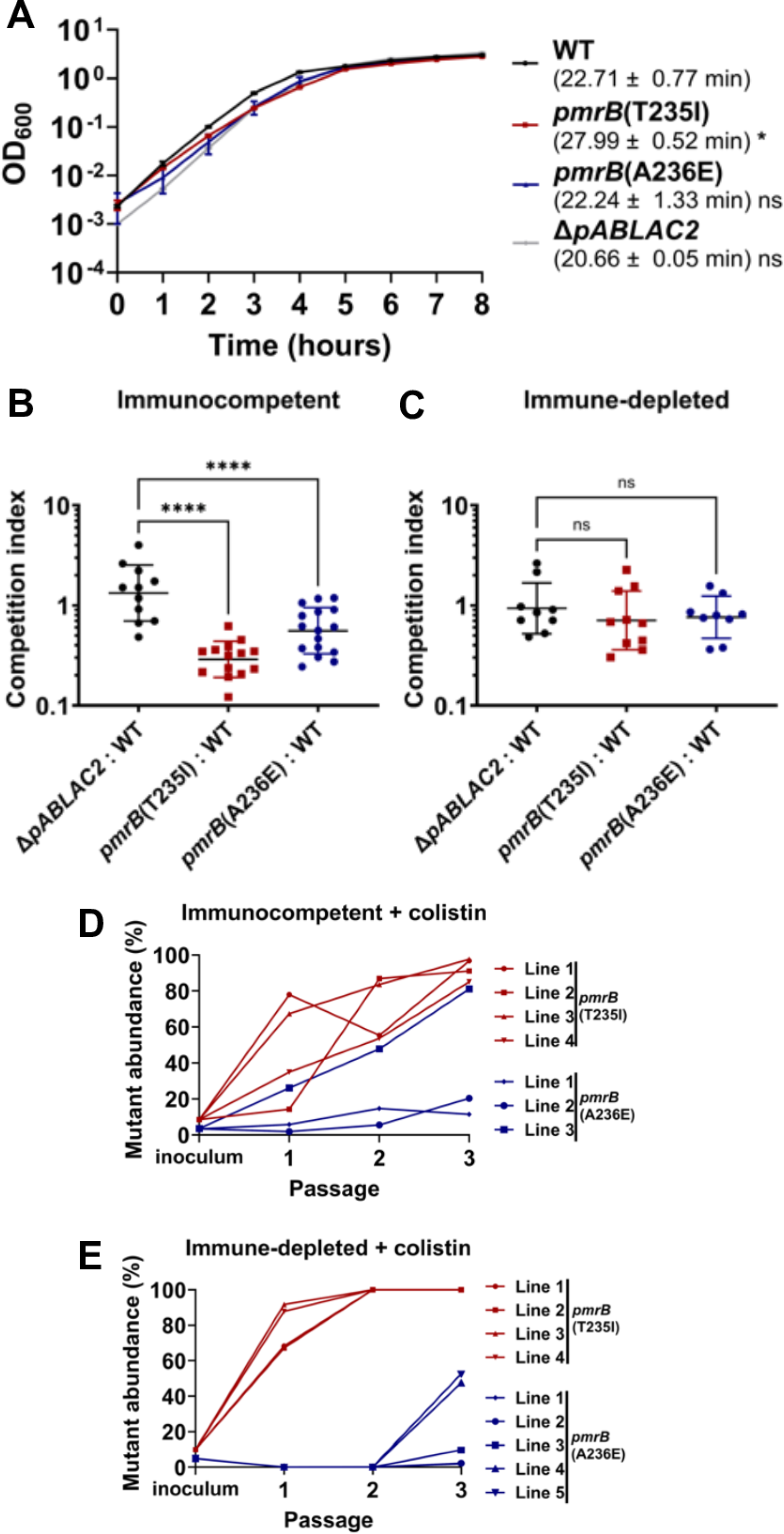
Mutations in *pmrB* selected during pulmonary infection incur a fitness cost in the absence of colistin. **A.** Kinetics of *pmrB* mutant growth in broth. Denoted LAC-4 strains were incubated in LB broth, and mass increase was measured at one-hour intervals. Mean doubling time ± SEM of three biological replicates shown. **B-C.** Competition of *pmrB* mutants vs. WT during pulmonary infections. Denoted strains were mixed at approximately 1:1 and inoculated via oropharyngeal aspiration into animals treated under noted conditions. 24hpi, mice were euthanized, and lungs were removed, homogenized in cold PBS, and plated on LB plates with or without colistin. Competition Index (CI) determined as described (Materials and Methods). Geometric Mean CI ± SD of 9-16 biological replicates was determined. **D-E.** The *pmrB*(T235I) strain efficiently competes with WT LAC-4 during pulmonary infections in the presence of colistin. WT LAC-4 and noted mutants were mixed at approximately 95:5 ratio followed by oropharyngeal inoculation into mice. 8mg/kg of colistin was administered 4hpi. At 24 hpi, the mice were euthanized, lungs were removed, homogenized, and plated on LB agar plates (Materials and Methods). Frozen pools from each passage were serially diluted and spot plated in LB plates supplemented with 0, 2 or 8μg/mL colistin. Mutant abundance was calculated as described (Materials and Methods). **A-C.** Statistical analysis of doubling time and CIs was performed using One-way ANOVA, followed by Dunnett’s multiple comparisons (Materials and Methods). \**P* < 0.05, \*\*\*\**P* < 0.0001; ns, not significant.

To reconstruct the fitness advantage of *pmrB* mutants relative to the parental control in the presence of colistin, each mutant was mixed with WT at 5:95 ratio and passaged through either immune-depleted or immunocompetent mice treated with colistin (Figs. 2D, E). After only two successive passages in the lung, the *pmrB*(T235I) mutant lines reached 100% of the total population in the immune-depleted host (Fig. 2E). Notably, during colistin treatment, this mutant also reached a high proportion of the population (>85%) in all lines after three passages in the immunocompetent host (Fig. 2D), even though this mutant competed poorly with wild type in the absence of drug pressure (Fig. 2B). The *pmrB*(A236E) mutant also showed evidence for effective competition during passaging in both host conditions, but constituted smaller proportions of the populations than that observed for the *pmrB*(T235I) mutant, with competitions not uniform among all the lines (compare Figs. 2D and 2E). Therefore, the *pmrB*(T235I) mutant selected in the immune-depleted conditions established a colistin-resistant population more efficiently than the *pmrB*(A236E) strain, even in the presence of an intact immune response.

### Colistin-resistant LOS-deficient strains show poor fitness during murine pneumonia

As the colistin-resistant mutations identified in the murine pneumonia model resulted in altered regulation of the PmrAB control system but did not directly disrupt LOS biosynthesis (Fig. 3A), we asked whether LOS biosynthesis dysfunction was an available mutational pathway to achieve colistin resistance during LAC-4 infection. A subset of *A. baumannii* clinical isolates can acquire resistance to polymyxins through mutations causing defective LOS biosynthesis (LOS^(−)^)(28). To determine whether the parental LAC-4 strain could tolerate LOS^(−)^ mutations, we first selected for colistin-resistant LAC-4 isolates and then screened for collateral vancomycin sensitivity (Fig. 3B). LOS^(−)^ mutants have a compromised outer membrane resulting in susceptibility to antibiotics usually ineffective against Gram-negative bacteria(27). We identified one such isolate and whole genome sequencing revealed a single *lpxA* G142R (*lpxA*\*) mutation, predicted to disrupt LOS synthesis (Dataset S2). SDS-polyacrylamide gel analysis of protease-treated extracts showed that, in contrast to the *pmrB* mutants, the *lpxA*\*-containing strain showed no evidence of LOS comigrating with the parental WT LAC-4 strain (Fig. 3A).

**Figure 3:**
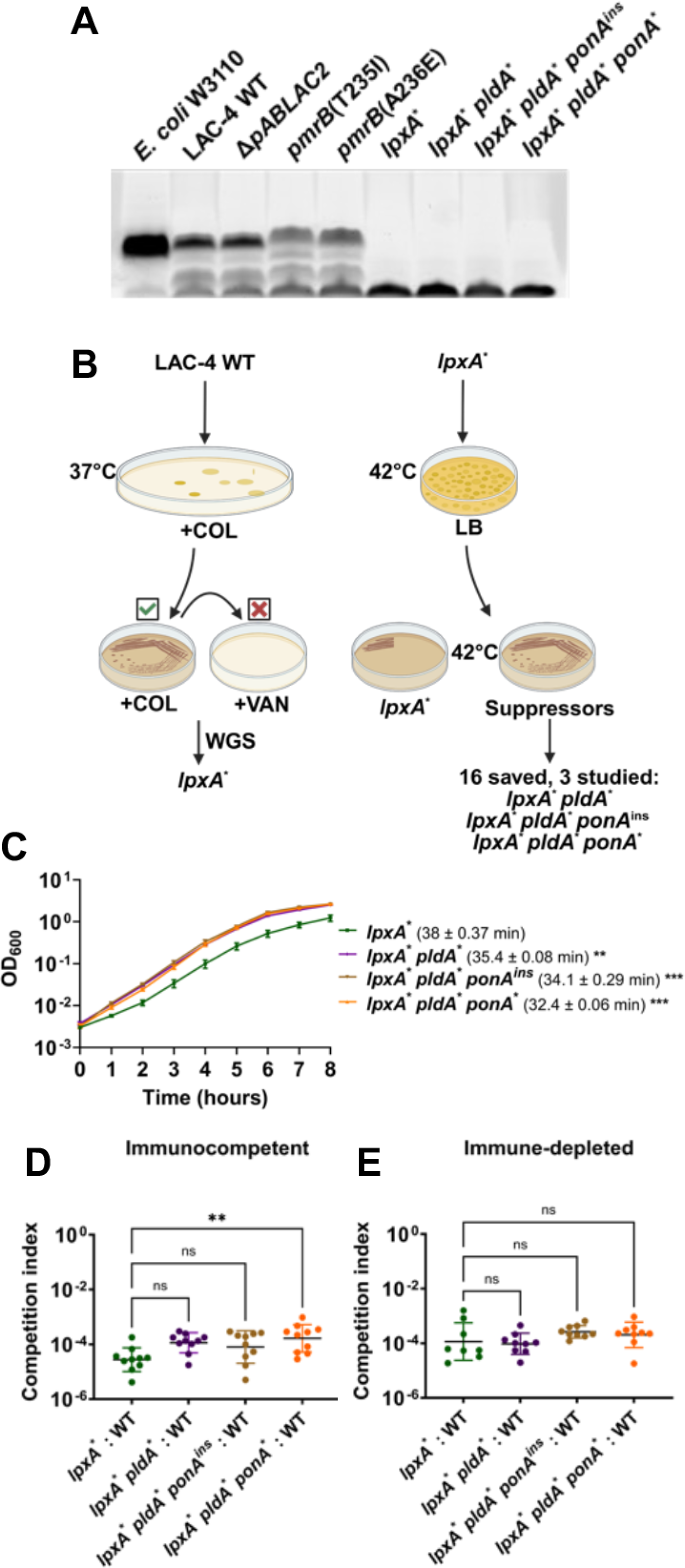
LOS-deficient *A. baumannii* isolates are defective for lung colonization. **A.** LOS^(−)^ strains lack carbohydrate sidechain. LOS isolated from denoted strains was fractionated on SDS-PAGE and carbohydrate was stained with ProQ Emerald 300. The denoted *E. coli* strain does not produce O-antigen and should migrate closely to *A. baumannii* LOS. The bottom band present in all samples is an unknown glycolipid or intermediate species. **B.** Protocol for selection of LOS-deficient LAC-4 and higher fitness variants. WT LAC-4 was plated on LB plates supplemented with 10μg/mL colistin and colonies were restreaked onto identical medium. An isolate (*lpxA*\*) that grew at 10μg/mL colistin but not 10μg/mL vancomycin was saved and sequenced. 10^9^ CFU of the isolate were plated on LB at 42°C and papillae growing above the lawn after 72 hrs. were streaked onto LB. Candidates showing improved growth at 42°C relative to parental LOS^(−)^ strain were saved and sequenced (Dataset S2). Three non-identical isolates (*lpxA*\* *pldA*\*, *lpxA*\* *pldA*\* *pbp1A*^ins^, and *lpxA*\* *pldA*\* *pbp1A*\*) were selected for further study. **C.** Kinetics of LOS^(−)^ mutant growth in broth. Denoted LAC-4 strains were incubated in LB broth, and mass increase was measured at one-hour intervals. Mean doubling time ± SEM of replicates shown. **D-E.** Competition of LOS^(−)^ mutants vs. WT during pulmonary infection. Denoted strains were mixed approximately 1:1 and inoculated via oropharyngeal aspiration into mice. 24hpi, mice were euthanized, lungs were removed, homogenized in cold PBS, and incubated on LB solid medium in presence or absence of colistin. Geometric Mean CI ± SD of 8-9 replicates determined as described (Materials and Methods). **C-E.** Statistical analysis of doubling time and CI’s were performed using one-way Anova followed by Dunnett’s multiple comparison. \*\**P* < 0.01, \*\*\**P* < 0.001; ns, not significant.

Due to low fitness, LOS^(−)^ strains revert to LOS^(+)^ or acquire second-site mutations that alter the glycerophospholipid composition of the outer membrane(27, 28, 32, 47). LOS^(−)^ suppressor mutants were selected by growing the LAC-4 *lpxA*\* mutant on LB solid medium at 42°C, a temperature that exacerbates its growth defect and results in a fine lawn of growth (Fig 3B). After selection, 16 non-reverted colonies that grew above the lawn were saved and subjected to WGS. Mutations identified in this fashion included a frameshift in *pldA,* as well as missense mutations in the *ponA* gene, both of which had been previously identified as increasing the growth rate of LOS^(−)^ strains (Dataset S2) (27, 32). A quadruple mutant containing a mutation in *msbA*, a gene involved in lipooligosaccharide transport across the membrane, was also isolated(48). Additionally, *pldA* mutants with second step mutations in ABLAC_25350, which encodes a hypothetical protein, were also isolated but were not studied further (Dataset S2).

Three non-identical LOS^(−)^ suppressor mutants were chosen for further analysis, each of which had either a *pldA* mutation or *pldA ponA* mutations (Fig. 3B). The three suppressor mutations showed no evidence of the reversion to LOS production, as the LOS profile of proteolyzed samples fractionated on SDS gels were indistinguishable from the *lpxA*\* mutant (Fig. 3A). Furthermore, analysis by thin layer chromatography (TLC) showed that these suppressor mutants remained LOS deficient (Fig. 4C). Growth rates in broth cultures were determined for each of the LOS^(−)^ strains. The *lpxA*\* mutant had a drastic increase in doubling time (38mins) compared to its WT parent (22mins) (compare Figs. 2A and 3C). All three LOS deficient suppressor mutants exhibited significant improvements in doubling time (32-35mins; Fig. 3C). Therefore LAC-4 tolerates LOS deficiency, and similarly to other *A. baumannii* strains, mutations in the *pldA* and *ponA* genes enhance fitness.

**Figure 4:**
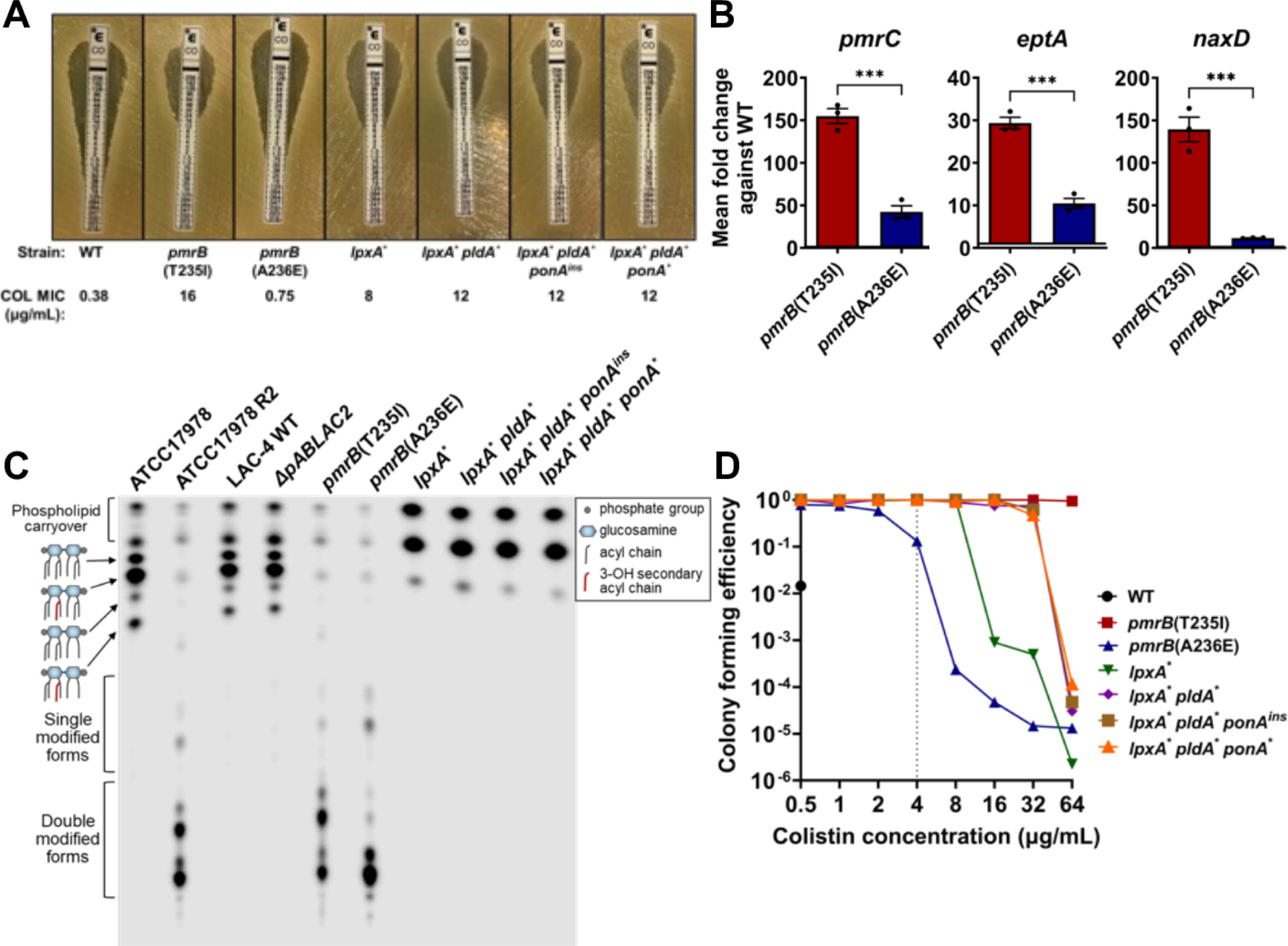
Overexpression of *pmrB*-regulated genes leads to colistin resistance and heteroresistance. **A.** Susceptibility of *pmrB* and LOS^−^ mutants to colistin. Colistin MIC determination via E-test strip. MIC determined as described (Materials and Methods). **B.** Expression of *pmrAB-*regulated genes in colistin-selected *pmrB* mutants. Transcript levels of *pmrC, eptA and naxD* was quantified via RT-qPCR and displayed as expression relative to WT. Mean ± SEM from three biological replicates shown. **C.** Lipid A from denoted strains was isolated, radiolabeled with ^32^P and separated by thin layer chromatography (TLC). Single and double modified forms refers to the addition of phosphoethanolamine and/or galactosamine. **D.** Population analysis profiling of *pmrB* and LOS^−^ mutants. Cultures of denoted strains were serially diluted and plated in LB supplemented with noted concentrations of colistin. Mean colony forming efficiency (CFE) was calculated as described (Materials and Methods). Grey vertical line: clinical breakpoint for resistance (≥4μg/mL). **B.** Statistical analysis of gene expression changes were performed using unpaired, two-tailed t-tests. \*\*\**P* < 0.001.

Although readily selected *in-vitro*, LOS^(−)^ clinical isolates have rarely, if ever, been observed and none emerged during passaging(49). Few studies have tested the virulence of LOS^(−)^ *Acinetobacter baumannii* mutants(50–52), and none, to our knowledge, have tested the impact of suppressor mutations on its pathogenesis. To determine their relative defects during murine pneumonia, the LOS^(−)^ strains were mixed at a 1:1 ratio with WT, inoculated into either immune-depleted or immunocompetent mice, and competitive indices were determined (Figs. 3D, E). In both host states, all LOS^(−)^ mutants had a devastating competition defect relative to WT, varying from 10^−4^ – 10^−5^ compared to the initial inoculum (Figs. 3D, E). This colonization efficiency was more than 1000X lower than observed for the *pmrB* mutants that had been selected during lung infection (compare Figs. 3D, E vs Figs. 2B, C). The suppressor mutations showed a fractional increase in the competitiveness relative to the parental *lpxA*\* strain when competitions were performed specifically in immunocompetent animals, with *lpxA*\* *pldA* ponA\**showing a significant increase in competitiveness (Fig. 3D). Even so, these suppressor mutations fail to compensate for the drastic colonization defects exhibited by LOS^(−)^ strains.

### Paths to colistin resistance occur via LOS modification and elimination

To both confirm and quantify the level of colistin resistance in our evolved strains, colistin E-test strip analysis was performed (Fig. 4A). All LOS^(−)^ strains surpassed the breakpoint for clinical resistance (≥4μg/mL) with the suppressor mutants showing colistin minimum inhibitory concentration (MIC) values that were higher than the *lpxA*\* parent (Fig. 4A). The WT LAC-4 strain was confirmed to be colistin-susceptible and the *pmrB*(T235I) mutant was highly colistin-resistant based on this MIC determination method (Fig. 4A). Surprisingly, the *pmrB*(A236E) mutant only had a slight increase in colistin MIC compared to WT based on the E-test determination and was clearly still susceptible based on the clinical drug resistance breakpoint definition (Fig. 4A).

As mutations in *pmrB* appear to be preferentially selected during disease because of their relatively high fitness compared to those lacking LOS(50–52), the genetic and biochemical basis of the two *pmrB* alleles was pursued. Similar to our observations modeling disease during mouse passage, colistin-resistant *Acinetobacter baumannii* clinical isolates primarily have mutations in *pmrB*(53). PmrB is the sensor kinase of the PmrAB two-component system whose regulon includes three genes involved in LOS modification(12). PmrC and its orphan homologue, EptA, are transmembrane proteins able to add phosphoethanolamine to both the 1- and 4’ phosphate groups in lipid A (SI Appendix, Fig. S2) (42). In addition, PmrAB positively regulates an operon including *naxD*, encoding a protein involved in the modification of the 1-phosphate group of lipid A via the addition of galactosamine (14).

To confirm that resistance in the *pmrB* mutants was due to modifications associated with upregulation of these genes, qRT-PCR analysis was performed. The transcript levels of *pmrC, eptA* and *naxD* were all increased in both *pmrB* mutants, with *pmrB*(T235I) resulting in higher upregulation than *pmrB*(A236E) relative to WT (Fig. 4B). When LOS preparations were analyzed by TLC, both mutants showed high levels of modified lipid A compared to WT, but the migration patterns of the modified forms were not identical in the two *pmrB* mutant strains (Fig. 4C). Serendipitously, the previously characterized colistin-resistant ATCC17978 R2 strain has a *pmrB*(T235I) mutation that results in lipid A modification as a consequence of both PmrC and NaxD activity(14, 42). The LAC-4 *pmrB*(T235I) mutant showed a similar lipid A profile to this strain, providing further evidence that colistin resistance in these mutants results from these two types of lipid A modification (Fig. 4C).

Mutations in *pmrB* in *A. baumannii* and other Gram-negative bacteria have also been associated with colistin heteroresistance, which can be confirmed by performing population analysis profiling (PAP)(54). This assay was performed with WT LAC-4 and the *pmrB* mutants, which revealed that *pmrB*(A236E) is heteroresistant due to its ability to form colonies at low frequencies at colistin concentrations 4-8X higher than the resistance breakpoint, even though the MIC is considerably lower than the breakpoint(18, 54) (Fig. 4D). In contrast, the WT was unable to grow at 1μg/mL colistin, which was 4-fold below the clinical breakpoint. The *pmrB*(T235I) mutant was distinct from both these strains, as it had 100% colony-forming efficiency up to 64μg/mL colistin (Fig. 4D). PAP of the LOS^(−)^ strains provided further evidence that the suppressor mutants have higher levels of colistin resistance compared to their *lpxA*\* parent (Fig. 4D). Therefore, the *pmrB* mutant selected in the immune-depleted host (T235I) was colistin-resistant while the one selected in the immunocompetent host (A236E) was colistin heteroresistant. Notably, in contrast to the high resistance strain, the heteroresistant mutant had a doubling time that was similar to WT during growth in broth culture (Fig. 2A).

### Insertion sequences drive heteroresistance in the *pmrB*(A236E) strain

Antibiotic heteroresistance is associated with strains that have subpopulations with higher levels of drug resistance relative to the parental strain(18). The high resistance exhibited in these subpopulations is often unstable and has been attributed to transient DNA amplification that allows hyperproduction of proteins that drive increased drug resistance(22). Consistent with this model, colistin heteroresistance in *Salmonella typhimurium* is driven by amplification of the *pmrD* gene, which encodes a positive regulator of *pmrAB*(24), and is consequently unstable via homologous recombination. *A. baumannii* does not encode *pmrD*, and little is known about what causes colistin heteroresistance in this organism other than it being linked to activating *pmrAB* mutations that drive upregulation of lipid A modifying enzymes(17). Therefore, starting with the low resistance *pmrB*(A236E) strain, we selected for increased resistance followed by continued growth in the absence of colistin selection to: 1) identify the genetic basis of variants that show high colistin-resistance in this strain; 2) test the stability of the resistant variants in the absence of colistin(18).

Three separate lines of the *pmrB*(A236E) strain were grown for 24 hours in LB broth supplemented with either 2μg/mL (Fig. 5A) or 4μg/mL (Fig. 5B) colistin, which were then passaged for 7 days in the absence of antibiotics. After drug exposure, colony forming efficiency (CFE) at 16μg/mL colistin increased to close to 100% frequency from the initial ~5×10^−4^ CFE before drug exposure, demonstrating enrichment for the colistin-resistant subpopulation. Once colistin was removed, 5/6 tested lines retained their high CFE at 16μg/mL colistin after 7 days of passaging (Figs. 5A, B). These results demonstrate that colistin treatment selects for the resistant subpopulation derived from the *pmrB*(A236E) strain that largely remains stable in the absence of antibiotic.

**Figure 5:**
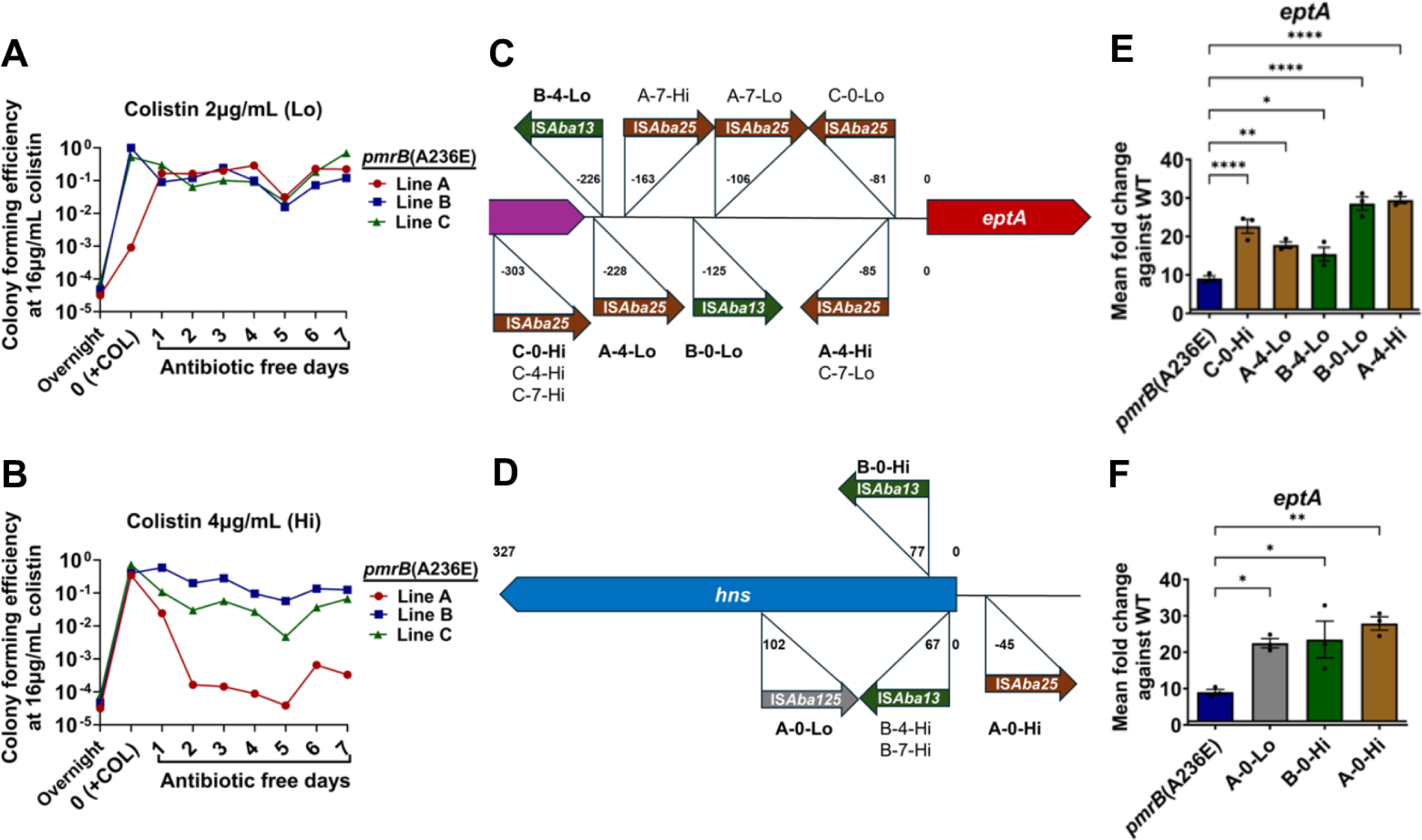
Enrichment of *pmrB*(A236E) isolates with higher colistin resistance selects for IS*Aba* activation of *eptA*. **A-B.** Enrichment and stability of high colistin resistance strains derived from the *pmrB*(A236E) background. Three lines (A, B, C) of the *pmrB*(A236E) mutant were selected for high resistance by 24 hr. growth in either 2μg/mL (Lo) or 4μg/mL (Hi) colistin (Day 0/+COL). Each line was subsequently passaged daily in antibiotic-free media for 7 days. Every 24 hrs., CFU were quantified and colony forming efficiency (CFE) on LB containing 16μg/mL colistin plates was determined, as described (Materials and Methods). Colonies growing at 16μg/mL colistin were isolated at day 0, 7, 14 for further analysis. **C-D.** Location and orientation of IS*Aba* insertions in colistin-enriched *pmrB*(A236E) isolates. Figures not drawn to scale. **E-F.** Expression of *eptA* in the colistin-enriched *pmrB*(A236E) isolates. Representative isolates (in bold) were selected for further analysis. Transcript levels of *eptA* in mutants relative to WT was quantified via RT-qPCR. Mean ± SEM from three biological replicates shown. **C-F.** Isolate nomenclature refers to “Line – Day collected – Level of colistin exposure at day 0”. **E-F.** Statistical analysis of gene expression changes were performed using one-way Anova followed by Dunnett’s multiple comparison. \**P* < 0.05, \*\**P* < 0.01, \*\*\*\**P* < 0.0001.

To detect genetic changes in the resistant *pmrB*(A236E) population, single colonies grown at 16μg/mL colistin were collected from all lines after 24 hours colistin treatment (day 0), and at 4 and 7 days after continuous passage in the absence of drug (Figs. 5A, B). E-test strips revealed that all 18 isolates had increased their colistin MIC compared to their *pmrB*(A236E) parent with 10/18 surpassing the breakpoint for clinical resistance (SI Appendix, Table S1, COL MIC). Short-read whole genome sequencing was performed on a single colony from each timepoint and confirmed they all retained the *pmrB*(A236E) mutation (SI Appendix, Table S1; Materials and Methods). Surprisingly, only 5/18 isolates contained additional mutations detected through short-read sequencing and only 1/18 exhibited evidence of genome amplification, which was a 9 gene region not known to be associated with antibiotic resistance (Dataset S3 and SI Appendix, Table S1).

Short-read sequencing analysis often fails to detect large insertions or other rearrangements in the genome(55). Long read sequencing and de-novo chromosomal assembly of the 18 isolates revealed that 17/18 isolates had acquired 1-4 insertion sequences (IS) at sites not found in the *pmrB*(A236E) parent (Dataset S3 and SI Appendix, Table S1). Long-read sequencing also revealed that the high resistance *pmrB*(T235I) strain contained the additional insertion mutations (ABLAC_16720::IS*Aba25*, ABLAC_23900::IS*Aba1*, and ABLAC_35150::IS*Aba13*) absent in the WT ancestor (Dataset S3 and SI Appendix, Fig. S3B).

The *A. baumannii* IS*Aba* insertion sequences are simple mobile genetic elements that can cause mutations by gene inactivation or altering expression patterns of transcripts downstream from transposition sites(56). The LAC-4 reference genome contains 6 different types of IS*Aba* inserted at 81 locations, comprising approximately 2.75% of the chromosome(36). IS*Aba* insertions into the region upstream of *eptA* and gene-inactivating insertions in *hns* have been observed in colistin-resistant *A. baumannii* isolates, both resulting in increased *eptA* transcription(15, 16). Consistent with insertions in these regions causing increased resistance, 16/18 of the colistin-selected *pmrB*(A236E) isolates contained IS*Aba* insertions upstream of *eptA* or in the *hns* region (Figs. 5C, D; Dataset S3; SI Appendix, Figs. S4C, D and Table S1). qRT-PCR analysis of *eptA* expression in representative isolates confirmed that IS*Aba* transposition at either site increased *eptA* expression but not that of *pmrC* or *naxD* (Figs. 5E, F; SI Appendix, Figs. S4C, D). Interestingly, several of the insertions upstream of *eptA* resulted in significantly lower expression of *naxD,* consistent with galactosamine modification of lipid A not being a critical determinant of increased colistin-resistance (SI Appendix, Fig. S4C)(12). TLC profiling of selected colistin-resistant isolates showed similar lipid A profiles compared to the heteroresistant parental *pmrB*(A236E) mutant (SI Appendix, Fig. S5). Thus, the stable colistin-resistant *pmrB*(A236E) subpopulation is predominantly created through IS*Aba* transposition events resulting in increased *eptA* expression.

### Insertion mutations remain stable in the absence of colistin treatment

The current models on heteroresistance postulate that the phenotypic stability of high drug resistance variants depends on the severity of the fitness cost of second-step mutations(18). The A-0-Hi isolate containing a single IS*Aba*25 insertion upstream of *hns* was chosen for further studies because it is the only second-step mutation identified in the *pmrB*(A236E) background that was associated with rapid loss of colistin resistance after subculture in the absence of drug (Fig. 5D, Dataset S3 and SI Appendix, Fig. S6A). qRT-PCR confirmed that the upstream insertion decreases *hns* gene expression in the A-0-Hi isolate compared to the *pmrB*(A236E) parent strain (*SI Appendix*, Fig. S6B). To test the stability of colistin resistance in this mutant, three separate lines of this A-0-Hi isolate were passaged for 14 days in the absence of antibiotic (Fig. 6A). Colony forming efficiency at 16μg/mL colistin remained stable for 9 days (~120 generations), then decreased to *pmrB*(A236E) levels between days 10-14 (Fig. 6A). Therefore, we were unable to reproduce the instability seen in the *pmrB*(A236E) Col Hi, line A after passaging in antibiotic-free media (Fig. 5B). However, *pmrB*(A236E) A-0-Hi was a single high-frequency isolate collected from this line and may not accurately represent the potentially mixed population that lost resistance.

**Figure 6:**
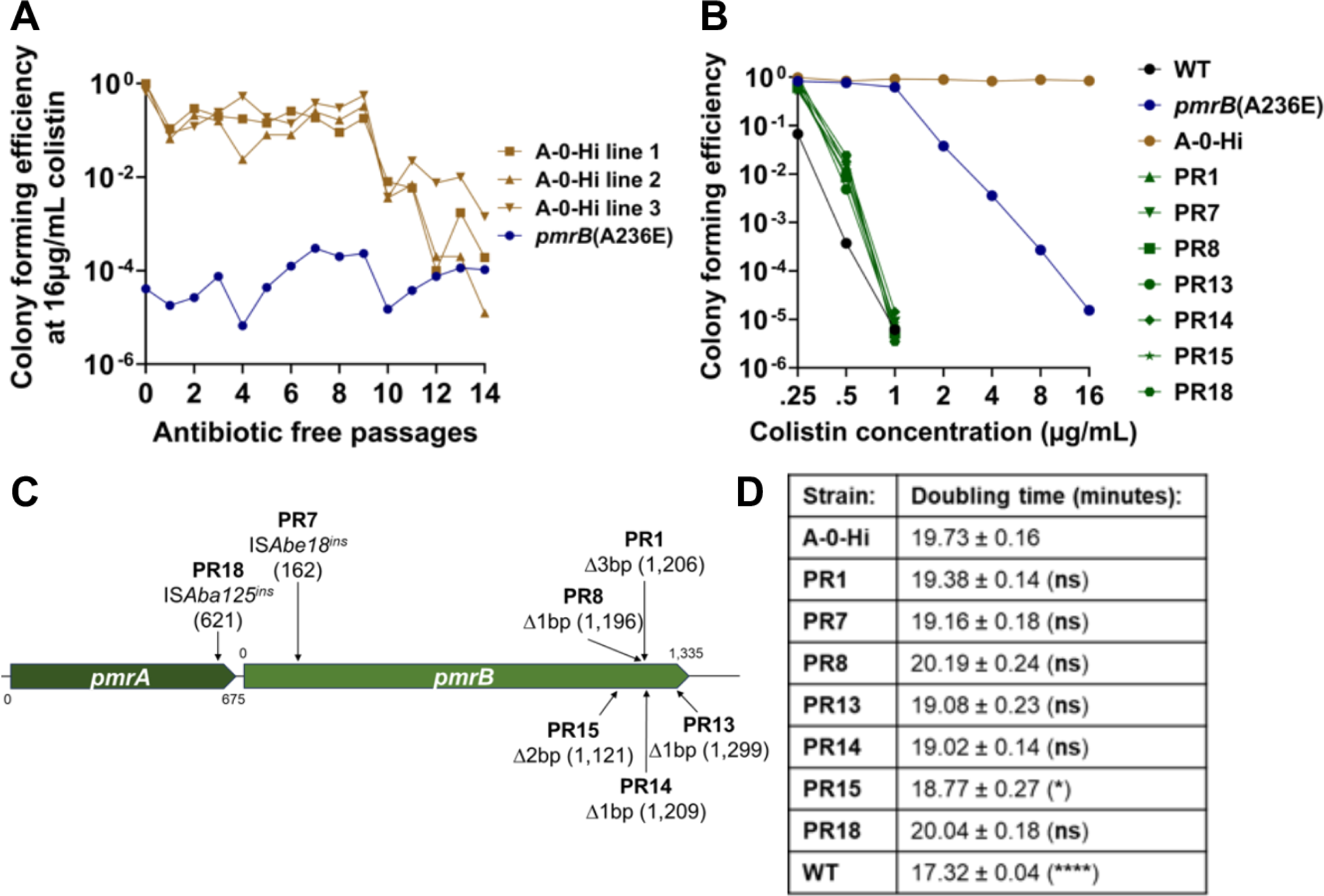
Colistin susceptible *pmrB*(A236E) A-0-Hi pseudorevertants emerge after continued passaging in the absence of drug. **A.** Stability of colistin resistance in the absence of antibiotic selection. Three lines of the colistin-resistant *pmrB*(A236E) A-0-Hi isolate were passaged in antibiotic-free media for 14 days. Every 24 hrs, colony forming units on LB plates containing 0 or 16μg/mL colistin was measured. CFE was calculated as described (Materials and Methods). **B-D.** 18 *pmrB*(A236E) A-0-Hi pseudorevertants (PR) were collected for further analysis by picking colonies growing on LB plates after 14 days of antibiotic-free passaging. **B.** Population analysis profiling of *pmrB*(A236E) A-0-Hi pseudorevertants. Cultures of denoted strains were serially diluted and plated on LB agar supplemented with noted concentrations of colistin. CFE was calculated as described (Materials and Methods). **C.** Schematic indicating identity and location of mutations in representative *pmrB*(A236E) A-0-Hi pseudorevertants detected by linear amplicon sequencing of *pmrCAB* region. **D.** Growth rates of *pmrB*(A236E) A-0-Hi pseudorevertants in broth. Mean doubling time + SEM of three biological replicates shown. Statistical analysis of doubling time was performed using One-way Anova followed by Dunnett’s multiple comparison. \**P* < 0.05, \*\*\*\**P* < 0.0001; ns, not significant. Kinetics shown in *SI Appendix,* Fig. S8A.

A-0-Hi isolates from all three lines were collected at day 14 from LB plates supplemented with 16 or 0 μg/mL colistin to identify the genetic changes that resulted in loss of resistance (Fig. 6A). Population analysis profiling confirmed that A-0-Hi colonies collected at day 14 from colistin-free plates, which we will refer to as pseudorevertants, had colistin sensitivity near WT levels and lower than their heteroresistant *pmrB*(A236E) ancestor (Fig. 6B). All A-0-Hi isolates collected at day 14 (either pseudorevertants or those remaining resistant) retained the IS*Aba25* insertion near *hns*, indicating that simple excision of the element was not the cause of the loss of drug resistance (*SI Appendix*, Fig. S7). Rather, linear amplicon sequencing of the *pmrCAB* region revealed that all 18 colistin-sensitive pseudorevertants acquired new mutations disrupting either the *pmrA* or *pmrB* genes (Fig. 6C and *SI Appendix*, Table S2).

To test whether reversion to sensitivity was due to the fitness cost incurred by high colistin resistance, growth rates in the absence of colistin were measured for the A-0-Hi strain and representative pseudorevertants (Fig. 6D and *SI Appendix*, Fig. S8A). Only one pseudorevertant exhibited a significant decrease in doubling time compared to its A-0-Hi parent, with none reaching WT growth rates (Fig. 6D). During the continuous passaging experiment, cultures were grown for hours in post-exponential phase prior to subculturing (Fig. 6A), so we reasoned that the pseudorevertants may have a competitive advantage after exponential growth. Therefore, we performed 1:1 competition experiments with all seven pseudorevertants against their A-0-Hi parent in LB broth and measured the competitive indices at 3 hours (mid-exponential) and 24 hours of culture (stationary). All pseudorevertants outcompeted their colistin-resistant parent at 24 hours, while none did so after 3 hours of culture (*SI Appendix*, Figs. S8B and C). Thus, colistin resistance in the *pmrB*(A236E) A-0-Hi insertion mutant is stable, but continuous passaging in the absence of drug eventually selects for colistin-sensitive pseudorevertants, with loss-of-function mutations in *pmrAB* being a common source of the selective advantage (Fig. 6C).

## Discussion

In the murine pneumonia model, immune depletion results in a permissive environment for *A. baumannii* replication, facilitating the emergence and selection of tolerant and drug-resistant mutants that fail to reach high numbers in immunocompetent animals(33). We provide further evidence that immune depletion accelerates the emergence of antibiotic resistance, with mutants having altered colistin sensitivity arising more readily in the immune-depleted passages. Furthermore, we found that bacterial fitness was another important determinant of resistance evolution during infection, as highly colistin-resistant mutants lacking LOS(49) lost the ability to colonize both immunocompetent and immune-depleted mouse lungs, even if they acquired secondary mutations that suppressed their growth defect in broth (Figs. 3C, E). The inability of LOS^(−)^ mutants to colonize the host, even after they acquire suppressor mutations, is heartening because LPS/LOS-targeting antibiotics that select for loss of LPS are likely to prevent proliferation during disease(57). Altogether, these findings highlight the role of the host tissue environment and the innate immune response in restricting the selection pool of antibiotic-resistant variants(46).

Regardless of the host state, colistin treatment during pulmonary disease selected for mutations that increased the expression of enzymes modifying LOS, with *pmrB* transcription-activating mutants being enriched in our model, as previously observed in broth culture models and in clinical isolates(58) (Figs. 1C, D). The evolved *pmrB* mutants were outcompeted by their drug-sensitive parent during lung infections of immunocompetent mice but not in immune-depleted hosts (Figs. 2B, C). Unlike the LOS^(−)^ mutants, both *pmrB* mutants retained the ability to grow in host tissues, allowing efficient competition with parental strains during colistin treatment (Figs. 2D, E). A previous study provided evidence that modified LOS from *A. baumannii pmrB* mutants induces higher TLR4 activation compared to unmodified LOS(59). This may be related to reports that some *pmrB* mutants have increased virulence, perhaps as a consequence of increased pattern recognition-driven inflammation(60, 61). Further work is needed to determine how LOS modifications caused by *pmrB* mutations impact the pathogenesis of *A. baumannii*.

We found that strains having mutations located one residue apart in the PmrB histidine kinase domain resulted in very different phenotypes (Fig. 4). Both mutants accumulated modified lipid A, with the doubly modified forms found at increased levels in the high resistance *pmrB(*T235I) strain compared to the heteroresistant *pmrB*(A236E) strain (Fig. 4C). This correlates well with the higher expression of *pmrC*, *eptA,* and *naxD* resulting from the *pmrB*(T235I) mutation (Fig. 4B). The high-resistance *pmrB*(T235I) mutant, however, was found to have three insertion element mutations that were absent both in the WT parent and the *pmrB*(A236E) strain (Dataset S3; COL17 vs. COL23), possibly explaining these resistance differences. When the *pmrB*(T235I) allele was backcrossed into WT LAC-4, the resulting strain had a colistin MIC (4μg/mL) that reached the clinical breakpoint for resistance, which is 8X higher than that of both the backcrossed and evolved *pmrB*(A236E) mutant strains (*SI Appendix*, Fig. S3A). Population analysis profiling showed that both the evolved and backcrossed *pmrB*(T235I) strains were able to form colonies all the way up to 64μg/mL colistin, with the evolved strain having a higher colony forming efficiency at this concentration as predicted by its higher E-test colistin MIC. In both backcrossed *pmrB* strains, an additional IS*Aba13* insertion was detected in identical locations and orientation 61bp upstream ABLAC_20540 (Dataset S3). Since this mutation does not confer resistance to the heteroresistant *pmrB*(A236E) backcrossed strain (*SI Appendix*, Figure S3), we can reasonably conclude that the *pmrB*(T235I) allele is responsible and sufficient for conferring LAC-4 with colistin resistance. Two additional IS*Aba* insertions mutations were detected in the backcrossed *pmrB*(A236E) strain (Dataset S3); underscoring the remarkable plasticity of the LAC-4 genome. These results indicate that *pmrB* mutations located one residue apart can lead to either colistin resistance or heteroresistance.

Although the resistance level of the *pmrB*(T235I) backcrossed strain was above the clinically defined breakpoint, it is notable that mouse passaging selected for a strain with even higher levels of resistance, possibly due to the three additional IS insertions (*SI Appendix*, Fig. S3A, COL17; MIC =16μg/mL). Evolved strains with the *pmrB*(T235I) mutation that harbored two of these insertion mutations (strains COL29-31) exhibited only a mildly increased MIC compared to the backcrossed mutant (*SI Appendix*, Fig. S3A) and did not outcompete the population during mouse passage (Dataset S1; *SI Appendix*, Figs. S1 and S3). The third mutation, found only in the highest resistance isolate, was in ABLAC_23900, encoding a putative TetR family regulator (*SI Appendix*, Fig. S3B). This triple insertion strain showed a 3X increase in *hns* expression compared to WT (*SI Appendix*, Fig. S6C), which could be attributable to any of these insertions, including one (ABLAC_35150::IS*Aba*13) located 513bp upstream of *hns* (*SI Appendix*, Fig. S6A). Coupled with our data showing that increased *eptA* expression is associated with insertions having decreased *hns* expression (Fig. 5E and *SI Appendix*, Fig. S6B), it is possible that misregulation of *hns* in any direction could increase colistin resistance via *eptA* overexpression, which is clearly a key step in the progression to colistin resistance.

A confounding factor in the colistin resistance literature is that mutations in *pmrB* have been associated with colistin susceptibility, resistance and heteroresistance(54). Indeed, our heteroresistant *pmrB*(A236E) mutation has been previously associated with colistin resistance(43). Based on the IS*Aba* insertions that we identified, the possibility emerges that colistin-resistant *pmrB*(A236E) strains may have secondary mutations that activate *eptA* expression. This is because the accumulation of added IS*Aba* insertions in the genome can be easily missed, even when performing whole-genome sequencing(55). This raises the possibility that heteroresistance-conferring mutations like *pmrB*(A236E) present a double danger, because their “susceptible” classification results in improper antibiotic treatment that is both toxic for the patient and selects for higher resistance mutants. We do not believe that the selection of these second-step insertion mutations is a phenomenon of growth in culture, as we were able to identify similar insertion mutations in *pmrB*(A236E) isolates after continued passage in the colistin-treated murine pneumonia model (Figs. 2D, E; *SI Appendix*, Fig. S9).

IS*Aba* transposition has been previously implicated in generating colistin-resistant isolates(19, 51, 59). It has been well-documented in several bacterial species that H-NS preferentially binds to chromosomal regions that are AT-rich relative to the rest of the genome(63). In *A. baumannii*, H-NS was shown to direct IS*Aba* transposition to these regions(62). Notably, the region upstream of *eptA* in LAC-4 is highly AT rich (79%) compared to the chromosome (61%), possibly explaining why 11/18 resistant isolates derived from *pmrB*(A236E) had IS*Aba* insertions in this region (Fig. 5C and Dataset S3) (16). In contrast, the *hns* gene itself, where 4/18 *pmrB*(A236E) high resistance isolates had insertions, is only 64% TA, but the upstream intergenic region is 80.9% TA (Dataset S3).

Most of the isolates derived from the heteroresistant *pmrB*(A236E) strain had IS*Aba* insertions in additional sites in the chromosome with unknown contributions to resistance (Dataset S3). One hotspot was in an intergenic region (ABLAC_36890-ABLAC_36900) that contained IS*Aba25* insertions in 6/18 *pmrB*(A236E)-derived isolates (*SI Appendix*, Fig. S10A). This region was also TA-rich (79%) and was flanked by genes involved in capsule synthesis (Dataset S3). Importantly, 5/6 of these isolates contained insertions in the vicinity of *eptA* and *hns* with the remaining isolate containing *eptA*(R127H) and *clpS*(Q90*) mutations that could contribute to colistin resistance (Figs. 5C, D; Dataset S3). The expression of the genes flanking these insertion mutations was tested via qPCR in two representative *pmrB*(A236E)-derived isolates, which contained either an *eptA-* or *hns*-disrupting insertion. In both isolates, the downstream gene ABLAC_36890 (annotated as *weeH*) had a significant (2X) reduction in expression while the upstream gene ABLAC_36900 only had increased expression when *hns* was disrupted (*SI Appendix*, Figs. S10B and C). Mutations upstream of *weeH* are associated with changes in the colony morphology, capsule, biofilm and motility of the mutant(62). The high frequency of transposition events observed in this study highlights the potential role of insertion elements in the adaptability of *A. baumannii*.

After ~120 generations of growth in broth, the colistin-resistant *pmrB*(A236) A-0-Hi strain was outcompeted by colistin-sensitive pseudorevertants (Fig. 6A, B). The majority of these pseudorevertants acquired loss-of-function mutations in *pmrB,* mainly in the region encoding the histidine-kinase like ATPase domain (Fig. 6C and *SI Appendix*, Table S2)(64). Other studies have observed colistin-resistant *A. baumannii* regain sensitivity after cessation of colistin therapy(11, 65). Previously characterized colistin-sensitive pseudorevertants with mutations disrupting the *pmrCAB* operon included isolates that may have been evolutionary dead ends, because they produced colistin-resistant mutants at significantly lower frequencies than their WT ancestors (65). There are several physiological changes in these revertants that could affect both fitness in culture and colonization in the host. Deletion of *pmrA* and *pmrB* in *A. baumannii* reduces motility, the production of biofilm, and the generation of outer membrane vesicles(66). In mouse models, transposon insertion mutations in *pmrAB* are colonization-proficient, indicating that loss-of-function of the regulator is compatible with maintenance of virulence (67, 68). It is not clear, however, if PmrAB inactivation provides an advantage during disease only when such isolates compete against strains that constitutively activate the system.

In summary, we have presented evidence that growth of the heteroresistant *pmrB*(A236E) parent in colistin selects for insertion mutations that drive acquisition of resistance (Figs. 5C, D; Dataset S3; *SI Appendix*, Table S1). Similar insertion mutations can be selected after colistin treatment of lungs infected with the *pmrB*(A236E) strain (Figs. 2D, E; *SI Appendix*, Fig. S9), or by simply incubating cultures of this heteroresistant mutant on agar plates supplemented with high colistin concentrations (*SI Appendix*, Fig. S11). Based on the results presented in this study, we propose the following model. Strains having heteroresistance mutations, such as *pmrB*(A236E), create a heterogenous population with varying degrees of lipid A modifications (Fig. 7A). This enables a subpopulation to survive colistin challenge, potentially creating a stress condition that activates transposition and results in resistance through insertions in the *eptA* or *hns* regions (Fig. 7B). After cessation of antibiotic treatment, colistin-sensitive pseudorevertants can emerge that acquire mutations disrupting *pmrAB* (Fig. 7C). Although this model predicts that the *pmrB*(A236E) isolates with resistance-conferring secondary mutations should show an increase in LOS modification relative to the *pmrB*(A236E) parent, this was not observed using the TLC strategy described here (*SI Appendix*, Fig. S5). This technique does not quantify the relative abundance of each type of single/modified lipid A form, such as double-modified phosphoethanolamine versus singly modified galactosamine plus phosphoethanolamine, which we suspect may underly increased colistin resistance (Fig. 7B).

**Figure 7.**
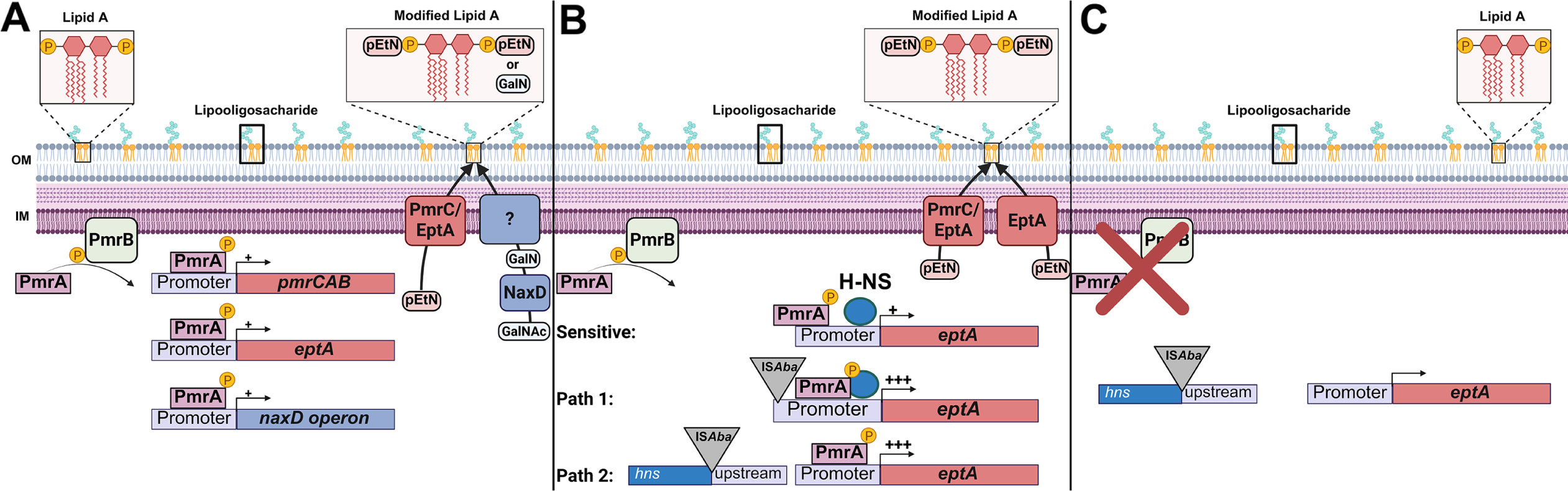
Model for the two-step acquisition of colistin resistance. **A.** Activation of the PmrB histidine kinase (via signal or mutation) leads to phosphorylation of PmrA. PmrA transcription factor binds to PmrA binding site in the *pmrCAB*, *eptA* and *naxD* promoter regions. PmrC and EptA add phosphoethanolamine (PEtN) to the 1 and 4’ phosphate groups on lipid A. NaxD deacetylates *N*-acetylgalactosamine (GalNAc) to galactosamine (GalN), which is then added to the 1-phosphate on lipid A by an unknown enzyme. These modifications reduce the negative charge of the lipid A portion of lipooligosaccharide which reduce its affinity to colistin. Mutations that strongly activate this system [ex. *pmrB*(T235I)] lead to resistance while mutations that weakly activate this system [ex. *pmrB*(A236E)] lead to heteroresistance. **B.** Heteroresistant *pmrB* mutant becomes fully colistin-resistant through insertion sequence transposition. The *eptA* gene is poorly activated by the *pmrB*(A236E) mutation, as the upstream region is TA rich and is silenced by H-NS (Sensitive). IS*Aba* transposition to the region upstream of *eptA* increases the expression of this gene leading to resistance (Path 1). IS*Aba* transposition into *hns* derepresses TA rich regions of the genome, including *eptA*, and leads to resistance (Path 2). **C.** Continued passaging in the absence of drug selects for colistin sensitive pseudorevertants that acquire loss of function mutations in *pmrAB*.

This work demonstrates that evolution of bacterial pathogens is a fine interplay between acquisition of antibiotic resistance and retention of fitness during growth and survival in tissues. As multidrug-resistant organisms continue to proliferate in healthcare and agricultural settings, accurately assessing the fitness costs of resistance mechanisms is crucial for identifying potential Achilles’ heels for novel treatments. Our work and others point to phosphoethanolamine transferases (ex. PmrC/EptA), which can spread through horizontal gene transfer in *mcr* plasmids(69), as attractive drug targets due to their involvement in colistin resistance and heteroresistance(70). Furthermore, IS transposition should be considered when studying antibiotic heteroresistance mechanisms that cannot be explained by gene amplification(22). Lastly, our results support the model that *A. baumannii* LOS^(−)^ mutants incur devastating fitness costs that prevent effective colonization during disease, highlighting the crucial role of this component in maintaining envelope integrity in Gram-negative bacteria(71).

## Materials and Methods

Unabridged Materials and Methods can be found in *SI Appendix*.

### Bacterial strains

All strains of *Acinetobacter baumannii* (AB) are derivates of LAC-4 or ATCC17978UN(38, 72). Bacteria were grown in Lysogeny Broth (LB) or LB agar plates at 37°C (*SI Appendix, SI Methods*).

### Molecular cloning and mutant construction

The *pmrB*(T235I) and *pmrB*(A236E) mutations were backcrossed from the COL17 and COL23 strains, respectively, into the parental WT LAC-4 background (SI Appendix, SI Methods). This was accomplished by electroporation and plasmid integration followed by homologous recombination, as described previously(73).

### Measuring growth rate and competitive index in broth culture

Overnight cultures of denoted strains were diluted 1:1000 in fresh LB broth in tubes. Cultures were grown at 37°C in a rotating roller set to 56rpm. OD_600_ was measured at t=0, and at one hour time intervals through 6-8 hrs. The averaged log_10_ values ± SEM over time were plotted. Mean doubling time and SEM of 2-3 biological replicates was calculated (*SI Appendix, SI Methods*).

For competition experiments in broth, overnight cultures of denoted strains were adjusted to the same OD, mixed at a 1:1 ratio and diluted into LB broth to a final OD of 0.002. Cultures were grown at 37°C for 3 and 24 hours. The number of colonies for the competing strains were quantified at hours 0, 3 and 24. Competition Index (CI) = output(Strain 1/ Strain 2) / input(Strain 1/ Strain 2). Geometric Mean CI and SD was plotted (*SI Appendix, SI Methods*).

### Selection of LOS deficient mutant and suppressors

Selection for a LOS deficient (LOS^(−)^) LAC-4 mutant was performed as described (32, 74) by screening for colistin-resistant colonies with collateral sensitivity to vancomycin (*SI Appendix, SI Methods*).

LOS^(−)^ LAC-4 mutants with suppressor mutations that improved growth rate were selected by plating the LOS^(−)^ strain in LB plates incubated at 42°C for 3 days (*SI Appendix, SI Methods*).

### Animal protocols

All animal procedures were approved by the Institutional Animal Care and Use Committee (IACUC) of Tufts University. All animals used in this study were 8-10 week-old female BALB/C mice obtained from Jackson Laboratories. All mice were housed in ventilated caging systems (10–15 air changes per hour) at temperatures of 68–79 °F (~20–26 °C) and 30–70% humidity, with 12 hr. light and dark cycle.

### Bacterial passaging in pulmonary infection model

All passaging procedures were performed in identical manner in both the immunocompetent and immune-depleted mice, with inocula adjusted as described in Results(33). Immune depletion was induced by cyclophosphamide pre-treatment. Lung infections were established by oropharyngeal aspiration(33, 75).. At 4hrs. post infection, 8mg/kg colistin was directly administered into the trachea using a liquid PenWu device (BioJane)(76). At 24-hours post infection, mice were sacrificed using CO_2_, and lungs were removed and processed. After outgrowing LAC-4 on LB plates, the bacteria were collected and frozen stocks were prepared (*SI Appendix, SI Methods*).

For passage 2-16, the frozen bacteria from the previous passage were revived and grown to mid-exponential phase at 37°C. These cultures were then adjusted to the appropriate OD_600_ and used to create inocula. The infection, drug administration, and bacterial harvest were performed in identical fashion at each passage (*SI Appendix, SI Methods*).

The same protocol was used for competitions between purified LAC-4 mutants and WT in the presence of colistin treatment in either immunocompetent or immune-depleted mice. To prepare the inocula, the mutants and WT were grown to mid-exponential phase, mixed at an approximate 5:95 ratio (mutant:WT), and inoculated into mice. Mutant abundance was calculated by quantifying number of colonies growing in drug plate (selective for *pmrB* mutants) over number of colonies in no drug plate and then multiplied by 100 (*SI Appendix, SI Methods*).

### Mouse competition assays

Mouse competitions were performed as described(33). Immune depletion was induced with cyclophosphamide pre-treatment and infections were initiated via oropharyngeal aspiration. At 24-hours post-infection, lungs were removed and homogenized aseptically in PBS. The number of colonies for the competing strains were quantified in the inocula and the lung homogenate. Geometric Mean CI and SD was plotted (*SI Appendix, SI Methods*).

### Isolation and quantification of mutants with altered colistin susceptibility

Pools from each passage of the pulmonary infection model were thawed and CFU were determined on LB plates in the absence of antibiotics and on graded concentrations of colistin (0 - 4μg/mL)(33). Colony forming efficiency (CFE) = number of colonies on colistin-containing plate/number of colonies in LB plate. For each colistin concentration tested, CFE was plotted as a function of passage number. Colonies growing at colistin concentrations not viable for WT were saved for further study (*SI Appendix, SI Methods*).

### Population analysis profiling (PAP)

Overnight cultures of WT LAC-4 and denoted mutants were diluted into fresh LB and grown to exponential phase. CFE were then determined on LB plates with increasing concentrations of colistin (0 - 64 μg/mL)(54). CFE were plotted as a function of colistin concentration for each strain (*SI Appendix, SI Methods*).

### Antibiotic MIC determination

Colistin MIC determinations were performed using the E-test strip method(54). MIC was determined by identifying the site where the zone of inhibition meets the strip (*SI Appendix, SI Methods*).

### Resistance stability assay

The resistance stability assay was performed as described(77). For the *pmrB*(A236E) passaging, three separate colonies were used to establish three lines. Overnight cultures for each line were grown in LB at 37°C. These cultures (input) were then diluted 1:100 in LB supplemented with either 2μg/mL (Lo) or 4μg/mL (Hi) colistin and grown for 24 hours at 37°C. After 24 hrs. drug exposure (Day 0), each culture was diluted 1:100 into fresh LB broth without antibiotic and was grown for 24 hours. Cultures were identically passaged in the absence of antibiotic for 7 days. At each day (input, day 0-7), colony forming efficiency at 16μg/mL colistin relative to absence of drug was determined. Isolates able to grow at 16μg/mL colistin were collected from day 0, 4, and 7 for further genome sequencing. For the *pmrB*(A236E) A-0-Hi passaging, three lines were passaged in the antibiotic-free media via 1:10,000 dilutions for 14 days. Isolates from day 14 growing at 0 and 16μg/mL colistin plates were saved (*SI Appendix, SI Methods*).

### Quantification of transcription by qPCR

Overnight cultures of denoted strains were diluted into fresh LB and grown at 37°C until mid-exponential phase. RNA was extracted using the Qiagen RNAeasy kit and subjected to cDNA synthesis using the Invitrogen SuperScript IV VILO kit. The qPCR reactions were performed using the Applied Biosystems PowerUp SYBR Green Master Mix and were run on a StepOnePlus Real-Time PCR system. Transcript level of target genes were normalized to 16S ribosomal RNA and fold change compared to WT was calculated using the ΔΔCt method (*SI Appendix, SI Methods*).

### Whole genome sequencing

Genomic DNA (gDNA) was purified from denoted isolates using the Qiagen DNeasy blood and tissue kit. Illumina Nextera reagents were used to prepare the libraries for short-read sequencing as described(33). 100bp single-end reads were obtained from pooled libraries using either a HiSeq2500 or a NovaSeq X Plus at the Tufts University Core Facility (http://tucf-genomics.tufts.edu/). Mutations were identified by applying the BRESEQ 0.38.8 pipeline to align the reads to the LAC-4 reference genome, including chromosome and two plasmids(78). (GenBank Accession: chromosome CP007712, pABLAC1 CP007713 and pABLAC2 CP007714)(36). See Dataset S4 for a comprehensive list of mutations detected in the evolved isolates.

For long-read sequencing, library preparation and sequencing was performed by the Hartwell Center for Biotechnology at St. Jude Children’s Hospital (SI Appendix). Alternatively, gDNA was sent to Plasmidsaurus for Nanopore long-read sequencing. Sequencing reads were de-novo assembled into genomes/contigs using the CANU pipeline(79). Mauve whole genome alignments on Geneious Prime were performed to compare the chromosomes of the mutants versus their parent. Insertions and deletions present in the mutants but not the parent were then identified (*SI Appendix, SI Methods*). All sequencing information provided in this text has been deposited in Genbank under the accession number PRJNA1273616.

### LOS and lipid A visualization

For fractionation and staining of whole cell LOS, 10^9^ CFUs of denoted strains were collected, washed and resuspended in 1X LDS sample buffer + 5% β-mercaptoethanol, and boiled for 10 minutes. Samples were then treated with proteinase K and run on a 4-12% Bis-Tris SDS-PAGE gel. Staining of LOS was performed using the ProQ Emerald 300 kit, following manufacturer instructions (80).

Lipid A isolation for TLC analysis from denoted AB strain was performed as described(80). Overnight cultures of denoted strains were diluted into fresh LB with 5 µCi/mL ^32^P ortho-phosphoric acid (Perkin-Elmer) and grown to an OD_600_ = 1. Lipid A was extracted using mild-acid hydrolysis followed by a Blight-Dryer extraction. Lipid A species were separated by thin layer chromatography (TLC) in a chloroform/pyridine/88% formic acid/water (50:50:16:5, vol/vol) solvent system. An Amersham Typhoon laser scanner with a phosphorimaging screen was used to image the dry plates.

### Statistical Analysis

GraphPad Prism was used for statistical analysis. For all statistical analysis, One Way ANOVA followed by Dunnet’s multiple comparison or unpaired two-tailed t-tests were employed. *P < 0.05, \*\**P* < 0.01, \*\*\**P* < 0.001, ****P < 0.0001; ns, not significant.

### Figure Design

Illustrations in Figures 1A, 3B and 7 were created in BioRender.com.

## Acknowledgements

This work is supported by NIAID U01AI124302, U19AI158076, U19AI142780 and R21AI128328. We would like to thank Albert Tai and Irina V. Grinvald for help with library preparations and the next-generation sequencing support provided by S10OD032203 via Tufts University Core Facility Genomics Core. We would like to thank the St. Jude Children’s Hospital Hartwell Center for Biotechnology for help with sequencing library preparation and Pacbio sequencing. We would like to thank Eddie Geisinger, Tim Van Opijnen and all the members of the Isberg lab for their feedback on the study.

## Author Contributions

J.H-B. and R.R.I devised the study. J.H-B., B.H., L.M.V., E.B.B. and G. I. C. T. performed the wet lab experiments. H.E. and J.M.R. performed long-red sequencing and provided consultation on library preparations. J.H-B and W.H. performed the sequencing analysis. L.M.V and M.S.T. performed the LOS analysis and provided analysis of results.

