## Supplementary Appendix: Supplementary Methods, Tables and Figures for "Resistance, heteroresistance and fitness costs drive colistin treatment failure during *Acinetobacter baumannii* pneumonia"

PNAS

**Author affiliations:**

**This PDF file includes:**

SI Methods
Figures S1 to S11
Tables S1 to S4
Legends for Datasets S1 to S4
SI References

**Other supporting materials for this manuscript include the following:**

Datasets S1 to S4

**SI Methods**

**Bacterial strains**. All strains of *Acinetobacter baumannii* (AB) are derivates of LAC-4 or ATCC17978UN(1, 2). Bacteria were grown in Lysogeny Broth (LB) or LB agar plates at 37°C. Broth cultures were grown in tubes and placed on a rotating roller drum set to 56rpm. Bacterial culture density was determined at 600nm ($OD_{600}$) using a cuvette and multiwell spectrophotometer (Biotek Synergy) to monitor growth. Frozen stocks were prepared to a final glycerol concentration of 25% and stored at -80°C.

**Molecular cloning and mutant construction.** The *pmrB*(T235I) and *pmrB*(A236E) mutations were backcrossed from the COL17 and COL23 strains, respectively, into the parental LAC-4 WT background. This was accomplished by electroporation and plasmid integration followed by homologous recombination, as described previously(3). The mutant alleles were amplified from COL17 and COL23 to generate fragments of 2999 bp and cloned into pJB4648APRA, a suicide plasmid encoding apramycin resistance ($Apra^R$) derived from pSR47S(4). The plasmids harboring the mutant alleles were then integrated *via* homologous recombination into *A. baumannii* LAC-4, selecting $Apra^R$ to generate heterozygotes. The resulting strains were then plated on LB+10% Sucrose at 30°C to select against the plasmid *sacB* allele, and screened for $Apra^S$. Recombinants that acquired the mutant *pmrB* allele were verified by sequencing amplified PCR products covering the *pmrB* region.

**Measuring growth rate and competitive index in broth culture.** Overnight cultures of denoted strains were diluted 1:1000 in fresh LB broth in tubes. Cultures were grown at 37°C in a rotating roller set to 56rpm. $OD_{600}$ was measured at t=0, and at one hour time intervals through 6-8 hrs. 2-3 biological replicates (cultures derived from distinct colonies), each the average of 2 technical replicates (biological replicate diluted into separate cultures), were performed per strain. The averaged $\log_{10}$ values ± SEM over time were plotted. Doubling time of each biological replicate was calculated using a previously published script(https://github.com/huoww07/calulate_bacteria_doubling_time) (5). Mean doubling time and SEM of 2-3 biological replicates was calculated (Fig. 2A, 3C and 6D).

For competition experiments in broth, overnight cultures of denoted strains were adjusted to the same OD, mixed at an approximate 1:1 ratio and diluted into LB broth to a final OD of 0.002. Cultures were grown at 37$^{o}$C in a rotating roller set to 56rpm. To calculate the ratio of the competing strains at denoted timepoints, aliquots of the cultures were 10X serially diluted and incubated on LB plates in presence or absence of colistin. The antibiotic concentration of the drug-supplemented plates was selected so that the colistin-resistant A-0-Hi strain, but not WT and the colistin sensitive pseudorevertants, were able to grow. 8µg/mL colistin was used for *pmrB*(A236E) A-0-Hi strains. Colistin resistance (ColR) strain CFU was determined by colonies on the antibiotic-containing plates, with Colistin sensitive (ColS) strain CFU = CFU on LB plate – CFU on antibiotic-containing plate. Competition Index (CI) = output(ColS/ColR) / input(ColS/ColR). Geometric Mean CI (each derived from individual cultures) and SD was plotted (Figs. S7A, B).

**Selection of LOS deficient mutant and suppressors**. Selection for a LOS deficient mutant was performed as described (6, 7). Overnight cultures of LAC-4 WT were diluted in LB and grown to mid-exponential phase. $10^9$ CFU were then spread on 150mm LB plates that were supplemented with 10µg/mL colistin sulfate (Sigma: PHR1605). Slow-growing colonies that emerged at days 2 and 3 post-inoculation were picked and streaked on colistin-containing plates (10µg/mL) and grown overnight at 37°C. Only one of the picked colonies was able to grow at colistin 10µg/mL. This isolate and a LAC-4 WT control were then streaked on LB plates in presence or absence of 10µg/mL vancomycin (Sigma: SBR00001), and the colistin-resistant isolate was unable to grow in presence of vancomycin, as predicted for having a defect in LOS biosynthesis.

Mutants that improve fitness of the LOS deficient ($LOS^{(-)}$) strain were identified in the following fashion(8). The $LOS^{(-)}$ strain was grown to exponential phase, and $10^9$ CFU was spread unto LB agar plates. These plates were incubated at 42°C for 3 days. At day 3, a lawn had formed, papillae were identified, streaked on LB plates and grown at 42°C overnight. The $LOS^{(-)}$ strain in the absence of temperature selection was streaked in parallel as a control. The following day, 18 of the purified papillae whose growth was visibly better than their $LOS^{(-)}$ parent were grown in LB and frozen at -80ºC in 25% glycerol for downstream analysis.

**Animal protocols.** All animal procedures were approved by the Institutional Animal Care and Use Committee (IACUC) of Tufts University. All animals used in this study were 8-10 week-old female BALB/C mice obtained from Jackson Laboratories. All mice were housed in ventilated caging systems (10–15 air changes per hour) at temperatures of 68–79 °F (~20–26 °C) and 30–70% humidity, with 12 hr. light and dark cycle.

**Bacterial passaging in pulmonary infection model.** All passaging procedures were performed in identical manner in both the immunocompetent and immune-depleted mice, with inocula adjusted as described in Results(5). Immune depletion was induced by the intraperitoneal injection of 150mg/kg and 100mg/kg of cyclophosphamide monohydrate (Sigma: C7397)  at days 4 and 1 prior to infection respectively(9). Lung infections were established by transiently anesthetizing mice through isoflurane inhalation followed by administering bacteria grown to mid-exponential phase (~10^8 CFU for immunocompetent and ~5x10^7-5x10^6 CFU for immune-depleted mice) via oropharyngeal aspiration(5, 10). For the first passage, all three lines were derived from separate colonies and each line was maintained separately throughout the passaging. At 4hrs. post infection, mice were anesthetized with isoflurane and 8mg/kg colistin sulfate (Sigma: PHR1605) was directly administered into the trachea using a liquid PenWu device (BioJane)(11). At 24-hours post infection, mice were sacrificed using $CO_2$, lungs were aseptically removed, then placed in ice-cold phosphate buffer saline (PBS) until they were processed with a tissue homogenizer (Omni). Lung bacterial burden (CFU/lung) was determined by performing 10-fold dilutions of lung homogenate on LB plates (Figs. 1B, C).

To outgrow LAC-4 (ciprofloxacin-resistant) and select against normal lung flora, the homogenate was plated on 150 mm LB agar plates containing 2μg/mL ciprofloxacin (Sigma: PHR1044) and incubated at 25°C overnight. The next day, the bacteria were scraped off the plates and resuspended in PBS to create 25% glycerol frozen stocks, or aliquoted into cryotubes stored at -80°C to be used for DNA purification.

For passage 2-16, the frozen bacteria from the previous passage were thawed, diluted in fresh LB, and grown to mid-exponential phase at 37°C. These cultures were then adjusted to the appropriate $OD_{600}$ and used to create inocula containing the same approximate number of bacteria, differing by immune state as described above. The infection, drug administration, and bacterial harvest were performed in identical fashion at each passage (Fig. 1A).

The same protocol was used for competitions between purified LAC-4 mutants and WT in the presence of colistin treatment in either immunocompetent or immune-depleted mice (Figs. 2D, E). The mutants and WT were grown to mid-exponential phase, mixed at an approximate 5:95 ratio (mutant:WT), and inoculated into mice. Colistin treatment and lung processing was performed as described above. After overnight growth, the bacteria were collected and saved for the next round of infections. Mutant abundance versus WT (colony forming efficiency; CFE) was determined by plating frozen stocks in LB plates with or without colistin. The colistin concentrations (2μg/mL for *pmrB*(A236E) and 8μg/mL for *pmrB*(T235I) of the drug-supplemented LB plates ensured only the mutants and not WT could grow. Mutant abundance was calculated by quantifying number of colonies growing in drug plate (selective for *pmrB* mutants) over number of colonies in no drug plate and then multiplied by 100. (Figs. 2D, E).

**Mouse competition assays.** Mouse competitions were performed as described(5). Immune depletion was induced with cyclophosphamide pre-treatment and infections were initiated via oropharyngeal aspiration (~$1X10^8$ CFU for immunocompetent group and ~$5x10^6$ CFU for immune-depleted group). At 24-hours post-infection, lungs were removed and homogenized aseptically in PBS. To calculate the ratio of the mutants to WT in both the inocula and the lung homogenates, the bacteria were 10X serially diluted and incubated on LB plates in presence or absence of colistin. The antibiotic concentration of the drug-supplemented plates was selected so that the mutants, but not the WT, were able to grow. 2μg/mL colistin was used for *pmrB*(A236E) and the LOS-deficient mutants, 8μg/mL was used for the *pmrB*(T235I) mutant, and 10ug/mL gentamycin was used to select against the gentamycin susceptible Δ*pABLAC2* mutant (Figs. 2B and 6E). Mutant CFU was determined by colonies on the antibiotic-containing plates, with WT CFU = CFU on LB plate – CFU on antibiotic-containing plate. Competition Index (CI) = output(Mutant/WT) / input(Mutant/WT). Geometric Mean CI (each derived from individual mouse infections) and SD was plotted (Figs. 2B, C; 3D, E).

**Isolation and quantification of mutants with altered colistin susceptibility.** Pools from each passage of the pulmonary infection model (Fig. 1B) were thawed and CFU were determined on LB plates in the absence of antibiotics and on graded concentrations of colistin (0.5μg/mL, 1μg/mL, 2μg/mL, 4μg/mL)(5). Colony forming efficiency (CFE) = number of colonies on colistin-containing plate/number of colonies in LB plate. For each colistin concentration tested, CFE was plotted as a function of passage number (Figs. 1C; S1).

Isolates, including the mutants characterized in this study, were purified and saved using the following protocol. Colonies from pools growing at a colistin concentration above that tolerated by WT were picked and re-streaked in LB colistin plates of the selected concentration. After overnight growth at 37°C, purified colonies were grown in LB broth and saved in 25% glycerol for downstream analysis. Only isolates able to grow and form isolated colonies at the selected colistin concentration were analyzed (Dataset S1, Table S1).

**Population analysis profiling (PAP).** Overnight cultures of LAC-4 WT and denoted mutants were diluted into fresh LB and grown to exponential phase. CFE were then determined on LB plates with increasing concentrations of colistin (0.5, 1, 2, 4, 8, 16, 32, 64 µg/mL)(12). Mean CFE were plotted as a function of colistin concentration for each strain (Figs. 4C, 6B; Figs. S3C, S11B). Each PAP experiment was repeated twice and a representative data is shown.

**Antibiotic MIC determination.** Polymyxin class antibiotics bind to the Costar plates(13), so colistin MIC determinations were performed using the E-test strip method(12). Overnight cultures of WT and mutants were diluted into fresh LB broth, grown to exponential phase, and adjusted to an $OD_{600}$ = 0.1 in PBS (approximating 0.5X MacFarland standard). A sterile swab was used to apply the bacteria to LB plates and colistin E-strips (Biomerieux) were placed on top. Plates were incubated overnight at 37°C and MIC was determined by identifying the site where the zone of inhibition meets the strip (Figs. 3A, S3A, Table S1, Dataset S1).

**Resistance stability assay.** The resistance stability assay was performed as described(14). For the *pmrB*(A236E) passaging, three separate colonies were used to establish three lines. Overnight cultures for each line were grown in LB at 37°C. These cultures (input) were then diluted 1:100 in LB supplemented with either 2µg/mL (Lo) or 4µg/mL (Hi) colistin and grown for 24 hours at 37°C. After 24 hrs. drug exposure (Day 0), each culture was diluted 1:100 (Fig. 5A, B) into fresh LB broth without antibiotic and was grown for 24 hours. Cultures were identically passaged in the absence of antibiotic for 7 days (Figs. 5A, B). At each day (input, day 0-7), colony forming efficiency at 16µg/mL colistin relative to absence of drug was determined (Figs. 5A, B). Isolates able to grow at 16µg/mL colistin were collected from day 0, 4, and 7 for further genome sequencing (Table 1).

For the *pmrB*(A236E) A-0-Hi passaging, three lines were passaged in the antibiotic-free LB via 1:10,000 dilutions for 14 days. Colony forming efficiency at 16µg/mL colistin relative to absence of drug was determined at each day (Fig. 6A). Isolates from day 14 growing at 0 and 16µg/mL colistin plates were saved.

**Quantification of transcription by qPCR**. Overnight cultures of denoted strains were diluted into fresh LB and grown at 37°C until mid-exponential phase. RNA was extracted following manufacturer's instructions (Qiagen RNAeasy: 74106) and subjected to cDNA synthesis using the SuperScript IV VILO kit (Invitrogen: 11756050). The qPCR reactions were performed using the PowerUp SYBR Green Master Mix (Applied Biosystems: A25742) and were run on a StepOnePlus Real-Time PCR system (Applied Biosystems) following manufacturer's instructions. Transcript level of target genes were normalized to 16S ribosomal RNA and fold change compared to WT was calculated using the ΔΔCt method. Three biological replicates, with 2 technical replicates each, were performed per strain. The values were plotted as mean $\pm$ SEM (Figs. 3C; 4E, F; S4C, D; S6B, C; S10B, C).

**Endpoint PCR.** Colonies from denoted strains were resuspended in ultra-pure water and lysed by a 5-minute incubation at 95°C. Colony PCR was performed using the OneTaq 2X master mix reagents (NEB: M0482S) on a Biorad T100 thermal cycler following manufacturer's instructions. Primers used are listed in Table S4. PCR products were visualized on a Biorad Gel Dock Go imaging system after running on a 1% agarose TAE gel (Figs. S3B, S7, S9; Figs. S11A, C). The DNA 1kb Plus Ladder (Invitrogen: 10787018) was used as a reference. PCR products were purified using the Qiaquick PCR purification kit (Qiagen: 28104). Denoted samples were sent to Plasmidsaurus for linear amplicon sequencing and mutations were detected by aligning sequences to the reference genome on Geneious Prime (Fig. 6C, Table S2).

**Whole genome sequencing.** Genomic DNA (gDNA) was purified from denoted isolates using the DNeasy blood and tissue kit (Qiagen: 69506). Illumina Nextera reagents were used to prepare the libraries for short-read sequencing as described(5). 100bp single-end reads were obtained from pooled libraries using either a HiSeq2500 or a NovaSeq X Plus at the Tufts University Core Facility (http://tucf-genomics.tufts.edu/). Mutations present above a 5% cutoff were identified by applying the BRESEQ 0.38.8 pipeline to align the reads to the LAC-4 reference genome, including chromosome and two plasmids(15). (GenBank Accession: chromosome CP007712, pABLAC1 CP007713 and pABLAC2 CP007714)(16). Mutations described in selected passaged strains are those not present in the ancestral LAC-4 WT strain. (Dataset S1, S2, S3). See Dataset S4 for a comprehensive list of mutations detected in the evolved isolates.

For long-read sequencing, library preparation and sequencing was performed by the Hartwell Center for Biotechnology at St. Jude Children's Hospital. gDNA samples were quantified using the Quant-It Pico Green dsDNA assay (Invitrogen) and quality was assessed by fragment size analysis using the Femto Pulse system with the Genomic DNA 165 kb kit (Agilent). Samples with a median size greater than 10 kb were sheared to 10 kb using Covaris g-Tubes in a volume of 150µl buffer (10 mM Tris pH 8.0, 0.1 mM EDTA). Four column passes at 6,000 rpm were performed (Eppendorf: 5424). Sheared DNA was concentrated by performing a 1:1 clean-up using SMRT beads (PacBio) and eluted in 46 µl of low EDTA TE. Samples with a median size of less than 10 kb were not sheared, and then brought up to 46 µl using the same buffer. Libraries were individually constructed from 0.3 - 1 ug of genomic DNA using the TPK 3.0 library preparation kit (PacBio), including the addition of barcoded PacBio SMRTbell adapters to allow for multiplexing.

Up to 96 libraries were pooled in equimolar amounts and loaded onto a single Revio SMRT cell at a concentration of 250 pM. Sequencing run conditions were performed as follows: 1) adaptive loading turned on (target of 0.85); 2) base kinetics turned on; 3) two hr. pre-extension time; 4) 30 hr. movie time. Alternatively, gDNA was sent to Plasmidsaurus for Nanopore long-read sequencing. Sequencing reads were de-novo assembled into genomes/contigs using the CANU pipeline(17). Mauve whole genome alignments on Geneious Prime were performed to compare the chromosomes of the mutants versus their parent. Insertions and deletions present in the mutants but not the parent were then identified (Dataset S3). All sequencing information provided in this text has been deposited in Genbank under the accession number PRJNA1273616.

**LOS and lipid A visualization**. For fractionation and staining of whole cell LOS, $10^9$ CFUs of denoted strains were collected, washed and resuspended in 1X LDS sample buffer + 5% β-mercaptoethanol, and boiled for 10 minutes. Samples were then treated with proteinase K and run on a 4-12% Bis-Tris SDS-PAGE gel. Staining of LOS was performed using the ProQ Emerald 300 (Thermo Fisher) kit, following manufacturer instructions (Fig. 3A) (18).

Lipid A isolation for TLC analysis from denoted AB strain was performed as described(18). Overnight cultures of denoted strains were diluted into fresh LB with 5 μCi/mL $^{32}P$ ortho-phosphoric acid (Perkin-Elmer) and grown to an $OD_{600} = 1$. Lipid A was extracted using mild-acid hydrolysis followed by a Blight-Dryer extraction. Lipid A species were separated by thin layer chromatography (TLC) in a chloroform/pyridine/88% formic acid/water (50:50:16:5, vol/vol) solvent system. An Amersham Typhoon laser scanner with a phosphorimaging screen was used to image the dry plates (Figs. 3A, S5).

**Statistical Analysis**. GraphPad Prism was used for statistical analysis. For all statistical analysis, One Way ANOVA followed by Dunnet's multiple comparison or unpaired two-tailed t-tests were employed. *$P < 0.05$, **$P < 0.01$, ***$P < 0.001$, ****$P < 0.0001$; ns, not significant.

**Figure Design.** Illustrations in Figures 1A, 3B and 7 were created in BioRender.com.

### SI Figures and Tables

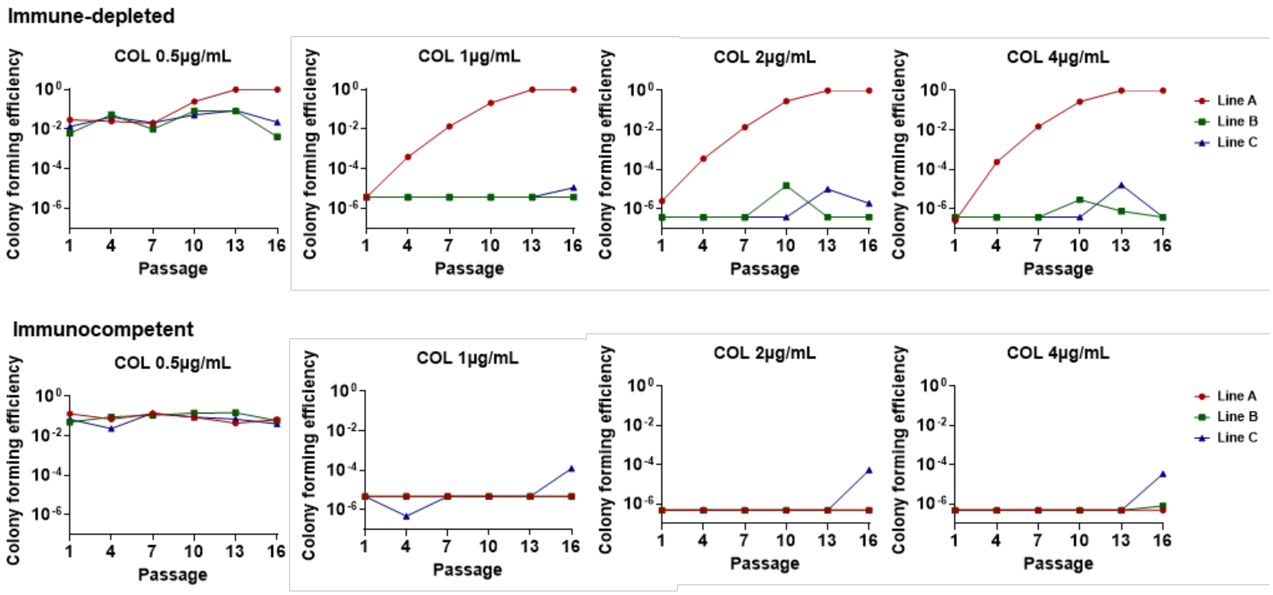


**Fig. S1. Quantification of mutants with altered colistin susceptibility in passaged pools.**

Frozen pools from each passage were incubated on LB agar containing 0-4 μg/mL colistin. CFE was calculated as described (Materials and Methods). Colonies appearing above baseline CFE were re-streaked on identical colistin-containing plates and saved for analysis (Dataset S1). The COL 4μg/mL results are identical to the data shown in Fig.1D-E and redisplayed here.

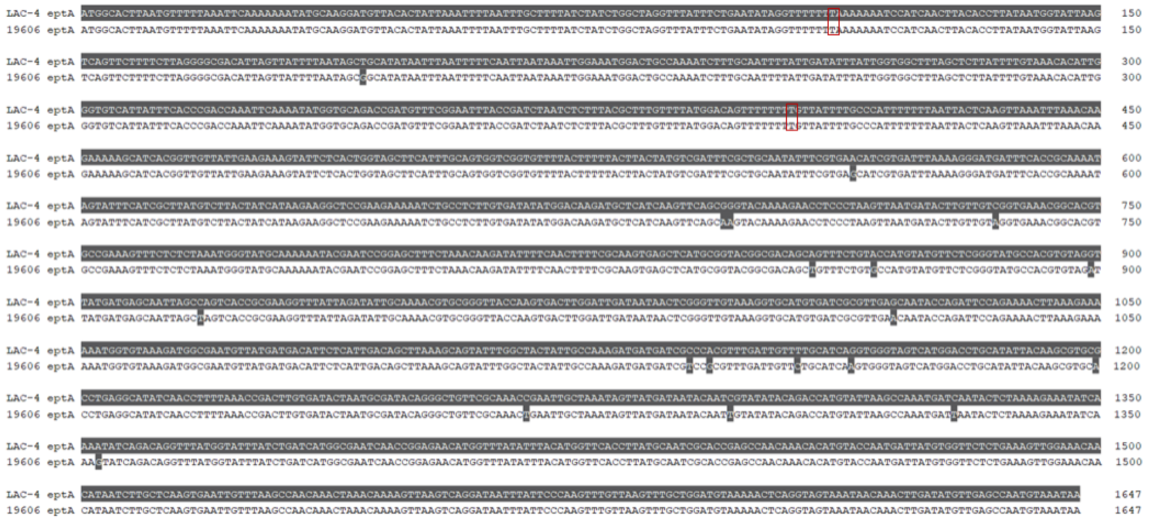

**Fig. S2. Alignment of LAC-4 and ATCC19606 *eptA*.**

The ATCC19606 *eptA* gene was mapped to the LAC-4 chromosome using Geneious Prime which shows 98.846% identity to this region. The LAC-4 reference genome had two missing T bases (red squares) at position 111 and 405/1647 causing this region to be annotated as two overlapping ORF's (ABLAC_18270-18280). WGS confirmed the presence of the T bases in the strains used in this study.

**Table S1: Colistin-enriched *pmrB*(A236E) isolates growing in 16µg/mL colistin plates.**

| Isolate: | Source: | COL MIC: (µg/mL) | Mutations: | IS*Aba* in vicinity of: | | | |
|---|---|---|---|---|---|---|---|
| | | | | *eptA* | *hns* | *weeH* | others |
| A-0-Lo | COL Lo, Line A, Day 0 | 6 | - | | ✓ | | |
| A-4-Lo | COL Lo, Line A, Day 4 | 1 | - | ✓ | | | |
| A-7-Lo | COL Lo, Line A, Day 7 | 2 | *gacS*(310C) | ✓ | | | |
| B-0-Lo | COL Lo, Line B, Day 0 | 2 | - | ✓ | | ✓ | ✓✓ |
| B-4-Lo | COL Lo, Line B, Day 4 | 1.5 | - | ✓ | | ✓ | ✓ |
| B-7-Lo | COL Lo, Line B, Day 7 | 6 | *clpS*(Q90*), *eptA*(R127H) | | | ✓ | |
| C-0-Lo | COL Lo, Line C, Day 0 | 3 | - | ✓ | | | |
| C-4-Lo | COL Lo, Line C, Day 4 | 4 | *clpA*(A302V) | | | | |
| C-7-Lo | COL Lo, Line C, Day 7 | 4 | - | ✓ | | | |
| A-0-Hi | COL Hi, Line A, Day 0 | 6 | - | | ✓ | | |
| A-4-Hi | COL Hi, Line A, Day 4 | 4 | - | ✓ | | | ✓✓ |
| A-7-Hi | COL Hi, Line A, Day 7 | 4 | - | ✓ | | | ✓✓ |
| B-0-Hi | COL Hi, Line B, Day 0 | 6 | - | | ✓ | ✓ | |
| B-4-Hi | COL Hi, Line B, Day 4 | 6 | *clpS*(+AC 132/414nt) | | ✓ | ✓ | |
| B-7-Hi | COL Hi, Line B, Day 7 | 8 | - | | ✓ | ✓ | ✓ |
| C-0-Hi | COL Hi, Line C, Day 0 | 1.5 | - | ✓ | | | |
| C-4-Hi | COL Hi, Line C, Day 4 | 1.5 | - | ✓ | | | ✓ |
| C-7-Hi | COL Hi, Line C, Day 7 | 2 | ABLAC_02920 (+T -46nt upstream) | ✓ | | | ✓✓ |

Mutations identified in high colistin-resistance *pmrB*(A236E) isolates from Fig. 5A and B. After selection at 2µg/mL (Lo) and 4µg/mL (Hi) colistin (COL), one isolate per line that grew to saturation in LB containing 16µg/mL colistin was collected at day 0, 4 and 7 and saved for whole genome sequencing (Figs. 5A, B). Isolate nomenclature refers to "Line – Day collected – Level of colistin exposure at day 0". Colistin MIC was determined by E-test strip. Mutations displayed were detected by short-read sequencing and all isolates retained their *pmrB*(A236E) mutation. IS*Aba* transposition into regions of interest was detected by performing long-read sequencing and assembling genomes *de-novo* (Materials and Methods). The vicinity of *eptA* refers to the region upstream this gene (ABLAC_18270-18280) including ABLAC_18260. The *hns* (ABLAC_35140) vicinity includes the gene itself and its upstream region. The *weeH* (ABLAC_36890) vicinity includes the intergenic region upstream this gene. See Dataset 3 for the other insertion mutations.

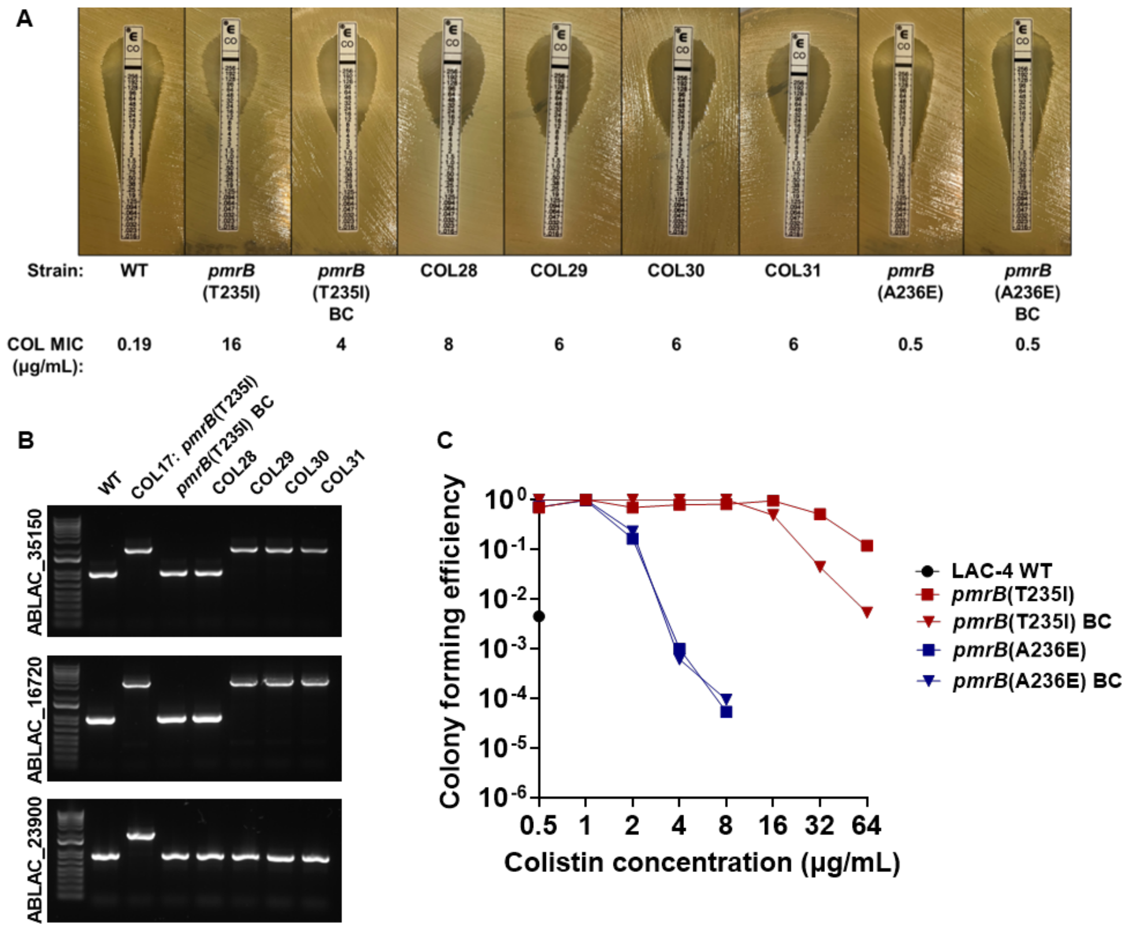


**Fig. S3. Colistin resistance is conferred by the *pmrB*(T235I) allele, with insertion mutations further augmenting the resistance observed in the evolved strains.**

**A.** Colistin susceptibility of isolates with *pmrB*(T235I) and *pmrB*(A236E) mutations. The backcrossed (BC) strains were generated via homologous recombination (Materials and Methods) and the COL28-31 isolates emerged during the lung passages (Dataset S1). Colistin MIC determined via E-test strip (Materials and Methods). **B.** PCR amplification of the denoted genes in isolates with the *pmrB*(T235I) mutation. WT was included as a negative (no insertion) control. See Dataset S3 for predicted function of these genes. **C**. Population analysis profiling of evolved and backcrossed *pmrB* mutants. Cultures of denoted strains were serially diluted and plated on LB agar supplemented with noted concentrations of colistin. CFE was calculated as described (Materials and Methods).

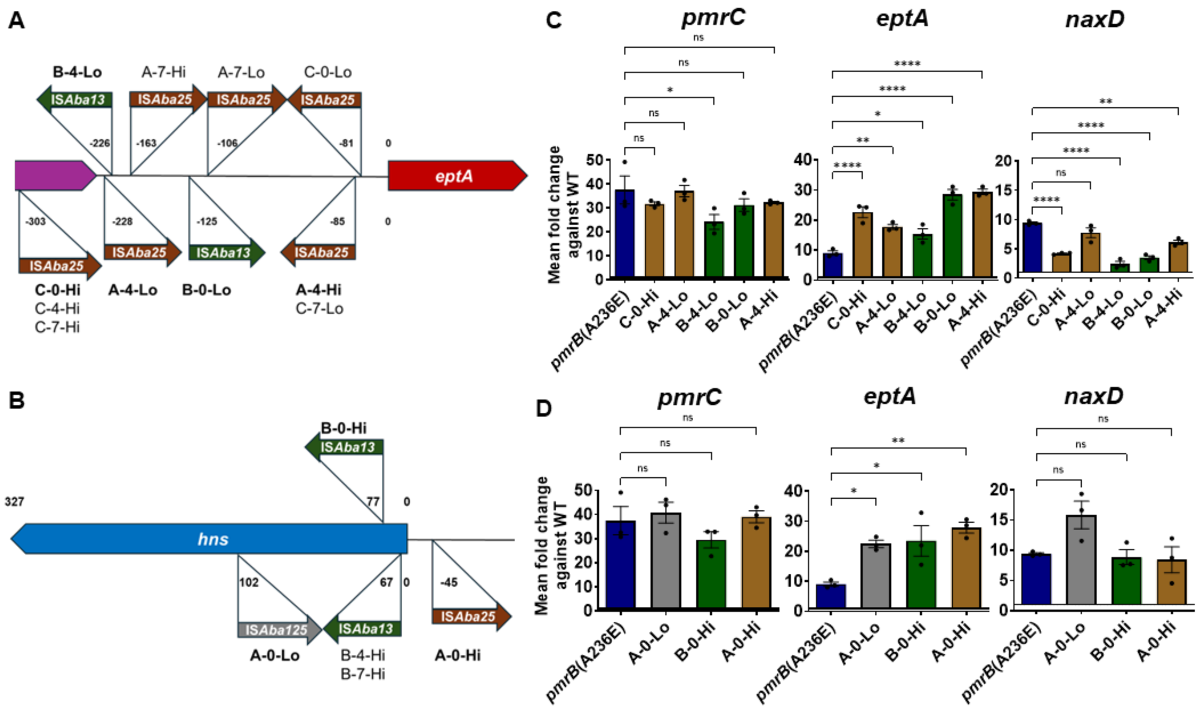


**Fig. S4. IS*Aba* insertions in the vicinity of *eptA* or *hns* do not increase *pmrC* or *naxD* expression.**

**A-B.** Location and orientation of IS*Aba* insertions in colistin-enriched *pmrB*(A236E) isolates. Figures not drawn to scale. **C-D.** Expression of genes that modify lipid A in the *pmrB*(A236E) isolates. Representative isolates (in bold) were selected for further analysis. Transcript levels of *pmrC, eptA and naxD* in mutants relative to WT was quantified via RT-qPCR. Displayed are mean $\pm$ SEM from three biological replicates shown. **A-D.** Isolate nomenclature refers to "Line – Day collected – Level of colistin exposure at day 0". **A-D.** The schematics and *eptA* data are the same shown in Figure 5. **C-D.** Statistical analysis of gene expression changes were performed using one-way Anova followed by Dunnett's multiple comparison. *$P < 0.05$, **$P < 0.01$, ****$P < 0.0001$.

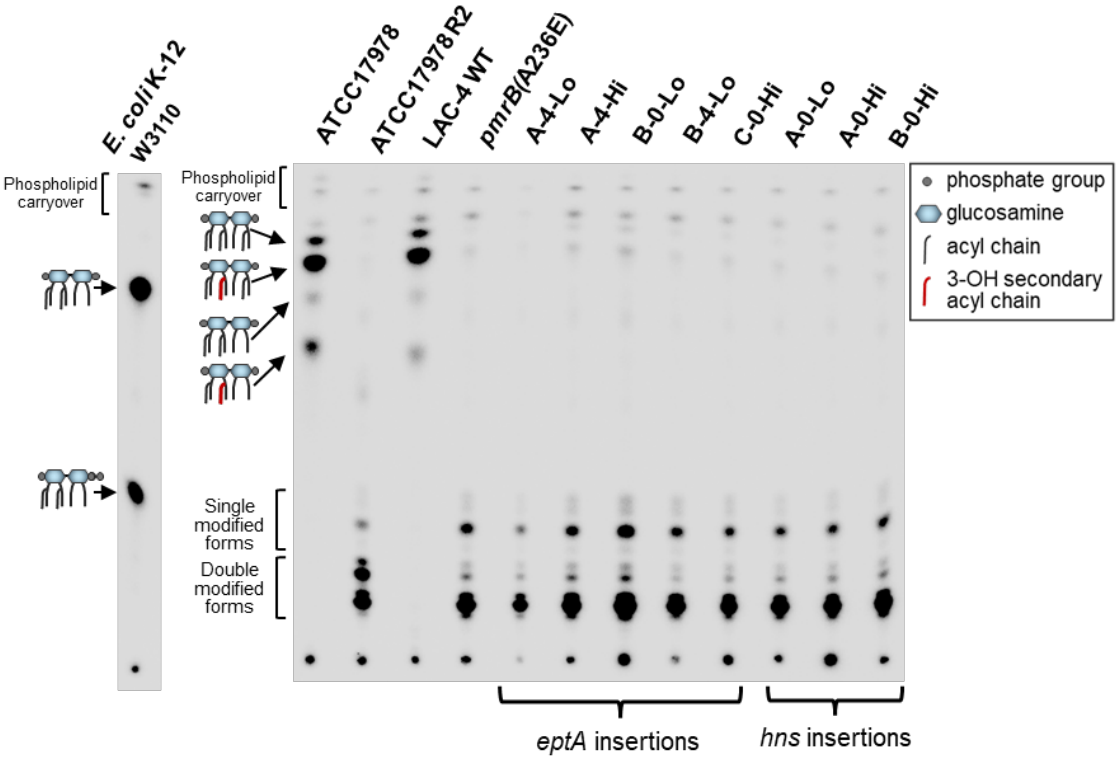


**Fig. S5. Lipid A profile of representative *pmrB*(A236E) isolates with insertions in the vicinity of *eptA* and *hns*.**

Lipid A profile of *pmrB* mutants. Lipid A from denoted strains was isolated, radiolabeled with $^{32}P$ and fractionated by TLC (Materials and Methods). Double-modified forms refer to the addition of phosphoethanolamine and/or galactosamine.

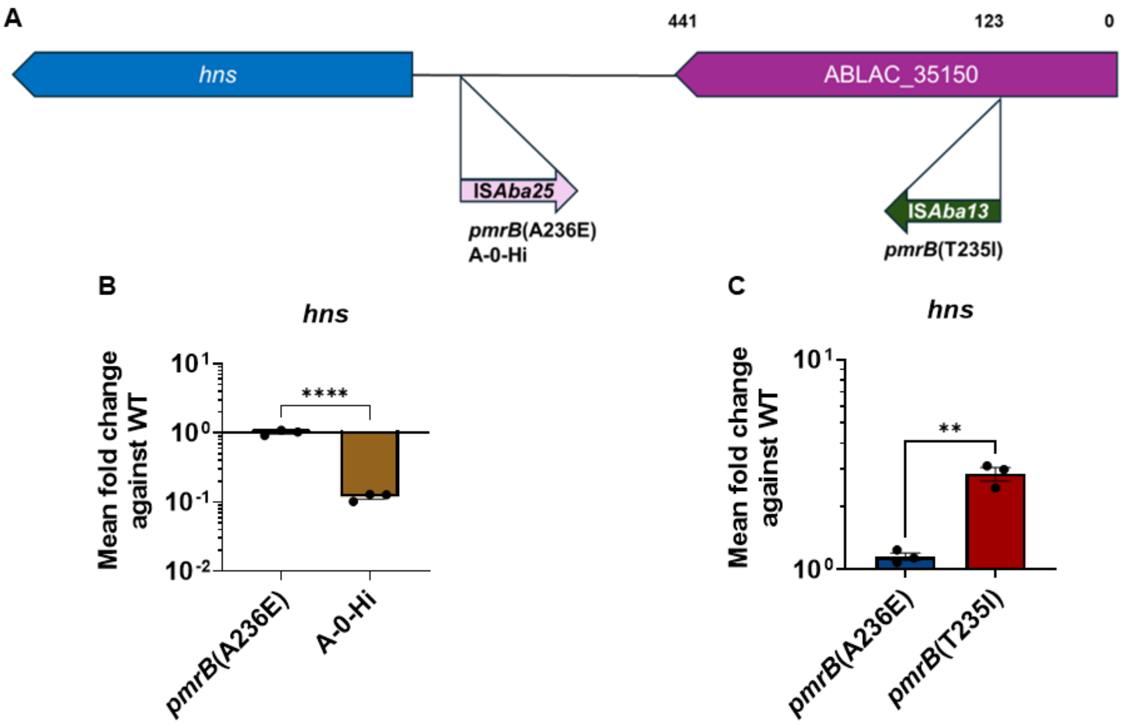


**Fig. S6. IS*Aba* insertions upstream *hns* alters this gene's expression.**

**A.** Location and orientation of IS*Aba* insertions upstream ABLAC_35140 (*hns*) in the *pmrB*(T235I) and the *pmrB*(A236E) A-0-Hi strains. Not drawn to scale. **B-C.** Expression of *hns* in denoted strains. Transcript levels of *hns* in mutants relative to WT were quantified via RT-qPCR. Mean ± SEM from three biological replicates shown. Statistical analysis of gene expression changes was performed using unpaired two-tailed t-tests. $^{**}P < 0.01$, $^{****}P < 0.0001$.

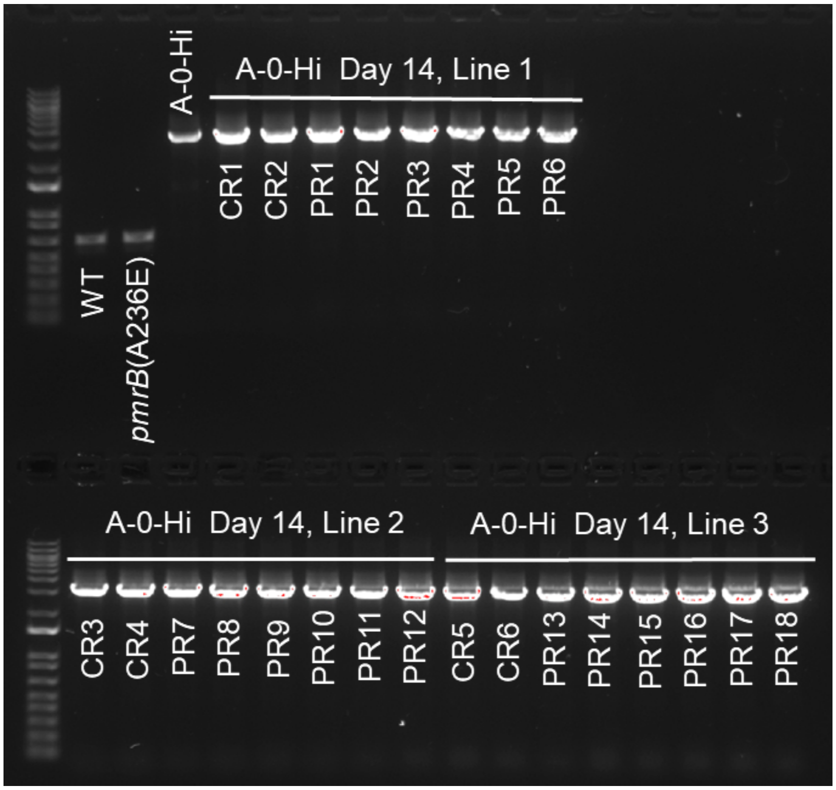


**Fig. S7. *pmrB*(A236E) A-0-Hi pseudorevertants retain their insertion upstream *hns*.** PCR amplification of the *hns* gene and its upstream region in denoted strains. After 14 days of antibiotic free passaging, 18 *pmrB*(A236E) A-0-Hi pseudorevertants (PR) were collected by picking colonies growing on LB plates and 6 colistin resistant (CR) colonies were collected from LB plates supplemented with 16µg/mL colistin (Fig. 6A). Colony PCR of the *hns* gene region was performed in denoted strains. The presence or absence of insertion mutations was visualized by running the PCR products on a 1% agarose TAE gel. WT and *pmrB*(A236E) were included as negative (no insertion) controls.

**Table S2: *pmrCAB* mutations present in *pmrB*(A236E) A-0-Hi pseudorevertants.**

| Isolate | Line | Gene | Mutation |
|---|---|---|---|
| **PR1** | 1 | *pmrB* | Δ3bp (1,206/1,335 nt) |
| PR2 | 1 | *pmrB* | Δ3bp (1,206/1,335 nt) |
| PR3 | 1 | *pmrB* | Δ3bp (1,206/1,335 nt) |
| PR4 | 1 | *pmrB* | Δ3bp (1,206/1,335 nt) |
| PR5 | 1 | *pmrB* | Δ3bp (1,206/1,335 nt) |
| PR6 | 1 | *pmrB* | Δ3bp (1,206/1,335 nt) |
| **PR7** | 2 | *pmrB* | IS*Abe18*$^{ins}$ (163/1,335 nt) |
| **PR8** | 2 | *pmrB* | Δ1bp (1,196/1,335 nt) |
| PR9 | 2 | *pmrB* | Δ1bp (1,196/1,335 nt) |
| PR10 | 2 | *pmrB* | Δ1bp (1,196/1,335 nt) |
| PR11 | 2 | *pmrB* | Δ1bp (1,196/1,335 nt) |
| PR12 | 2 | *pmrB* | Δ1bp (1,196/1,335 nt) |
| **PR13** | 3 | *pmrB* | Δ1bp (1,299/1,335 nt) |
| **PR14** | 3 | *pmrB* | Δ1bp (1,209/1,335 nt) |
| **PR15** | 3 | *pmrB* | Δ2bp (1,121/1,335 nt) |
| PR16 | 3 | *pmrB* | Δ1bp (1,299/1,335 nt) |
| PR17 | 3 | *pmrB* | Δ1bp (1,299/1,335 nt) |
| **PR18** | 3 | *pmrA* | IS*Aba25*$^{ins}$ (621/675 nt) |

Mutations identified in colistin-sensitive *pmrB*(A236E) A-0-Hi pseudorevertants (PR) from Fig. 6A. After 14 days of passaging in the absence of antibiotic, 6 high frequency isolates per line were collected from LB plates and saved for further analysis. Mutations displayed were detected by linear amplicon sequencing of the *pmrCAB* region (Materials and Methods). All isolates retained their *pmrB*(A236E) mutation. Isolates in bold were selected for further analysis and additional mutations detected via whole genome sequencing are listed in Dataset S4.

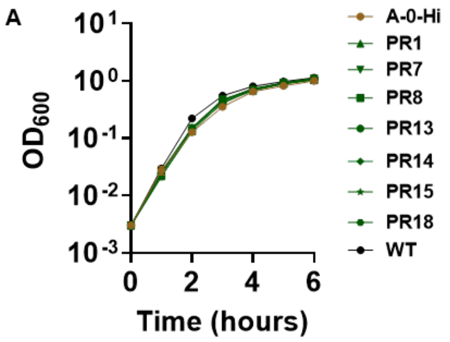


| Strain: | Doubling time (minutes): |
|---|---|
| A-0-Hi | 19.73 ± 0.16 |
| PR1 | 19.38 ± 0.14 (ns) |
| PR7 | 19.16 ± 0.18 (ns) |
| PR8 | 20.19 ± 0.24 (ns) |
| PR13 | 19.08 ± 0.23 (ns) |
| PR14 | 19.02 ± 0.14 (ns) |
| PR15 | 18.77 ± 0.27 (*) |
| PR18 | 20.04 ± 0.18 (ns) |
| WT | 17.32 ± 0.04 (****) |

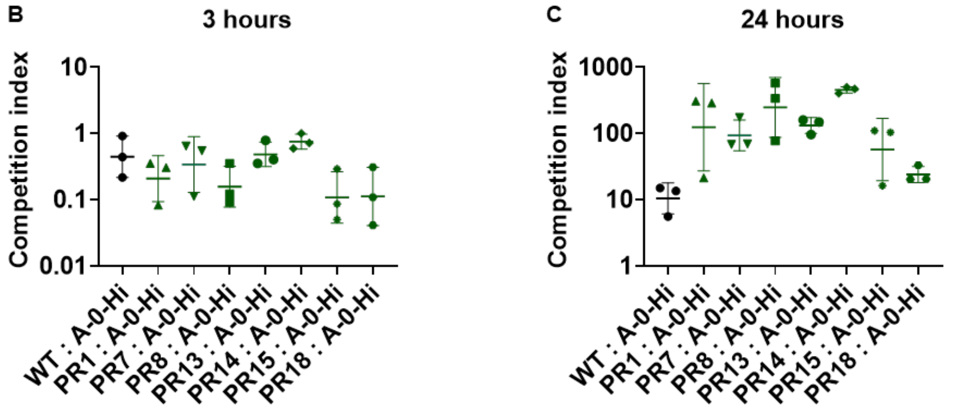


**Fig. S8. A-0-Hi pseudorevertants outcompete their resistant parent when grown past exponential phase.**

**A.** Kinetics of *pmrB*(A236E) A-0-Hi pseudorevertants growth in broth. Denoted LAC-4 strains were incubated in LB broth, and mass increase was measured at one-hour intervals. Mean doubling time + SEM of three biological replicates shown. Statistical analysis of doubling time was performed using One-way Anova followed by Dunnett's multiple comparison. *$P < 0.05$, ****$P < 0.0001$; ns, not significant. The table is a duplicate of Figure 6D. **B-C.** Competition of pseudorevertants versus their *pmrB*(A236E) A-0-Hi parent in broth. Denoted strains were mixed at approximately 1:1 and inoculated into LB broth. The ratio of the colistin sensitive strains to their resistant parent was measured at hours 0, 3 and 24. Competition Index (CI) determined as described. Geometric Mean CI + SD of 3 biological replicates shown.

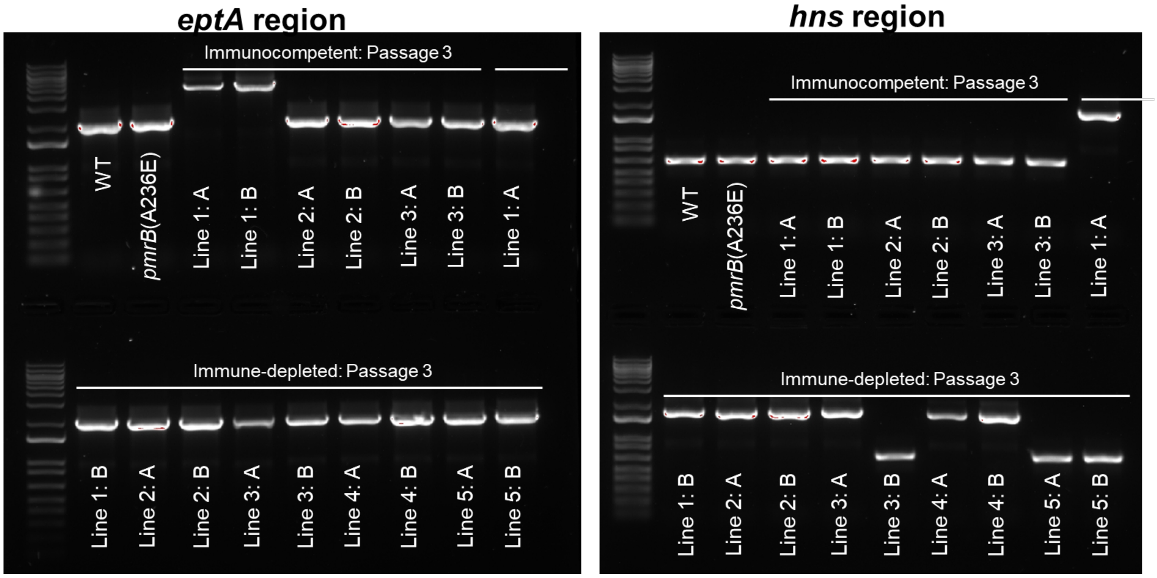


**Fig. S9. Insertions in the vicinity of *eptA* and *hns* can be detected in *pmrB*(A236E) isolates passaged in mice treated with colistin.**

Detection of insertions in the vicinity of *eptA* and *hns*. Frozen pools from passage 3 of the WT:*pmrB*(A236E) competitions in the presence of colistin treatment (Fig. 2D, E) were diluted and plated in LB plates supplemented with colistin 2µg/mL to select for *pmrB*(A236E) colonies and against WT. Two colonies per line (from both the immunocompetent and immune-depleted pools) growing at 2ug/mL colistin were re-streaked on identical colistin-containing plates. Colony PCR of both the *eptA* and *hns* gene regions was performed in denoted strains. The presence of new insertion mutations was visualized by running the PCR products on a 1% agarose TAE gel. WT and *pmrB*(A236E) were included as negative (no insertion) controls.

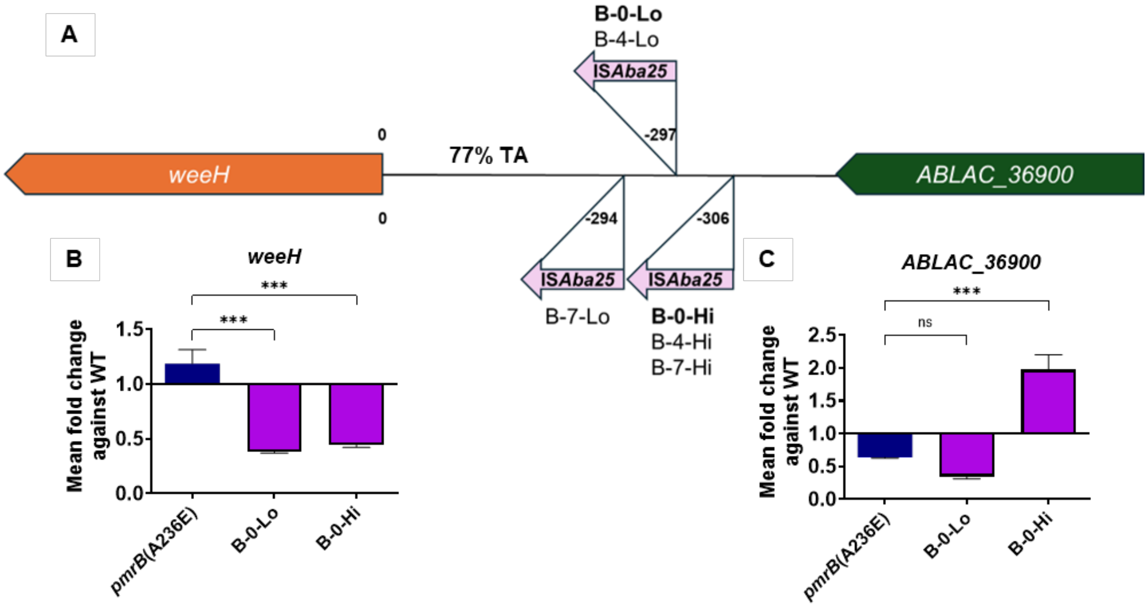


**Fig. S10. IS*Aba* insertions located upstream of *weeH* reduce expression of downstream gene.**

**A.** Location and orientation of IS*Aba* insertions upstream ABLAC_36890 (*weeH*) in colistin-enriched *pmrB*(A236E) isolates described in Figure 5. Representative isolates (in bold) were selected for further analysis. B-0-Lo contains an IS*Aba* insertion upstream *eptA* and B-0-Hi contains one in the *hns* gene. Not drawn to scale. **B-C.** Expression of genes flanking the IS*Aba* insertions upstream *weeH*. Transcript levels of *weeH* (ABLAC_36890) and ABLAC_36900 in mutants relative to WT was quantified via RT-qPCR. Mean $\pm$ SEM from three biological replicates shown**.** Statistical analysis of gene expression changes were performed using one-way Anova followed by Dunnett's multiple comparison.****P* < 0.001; ns, not significant.

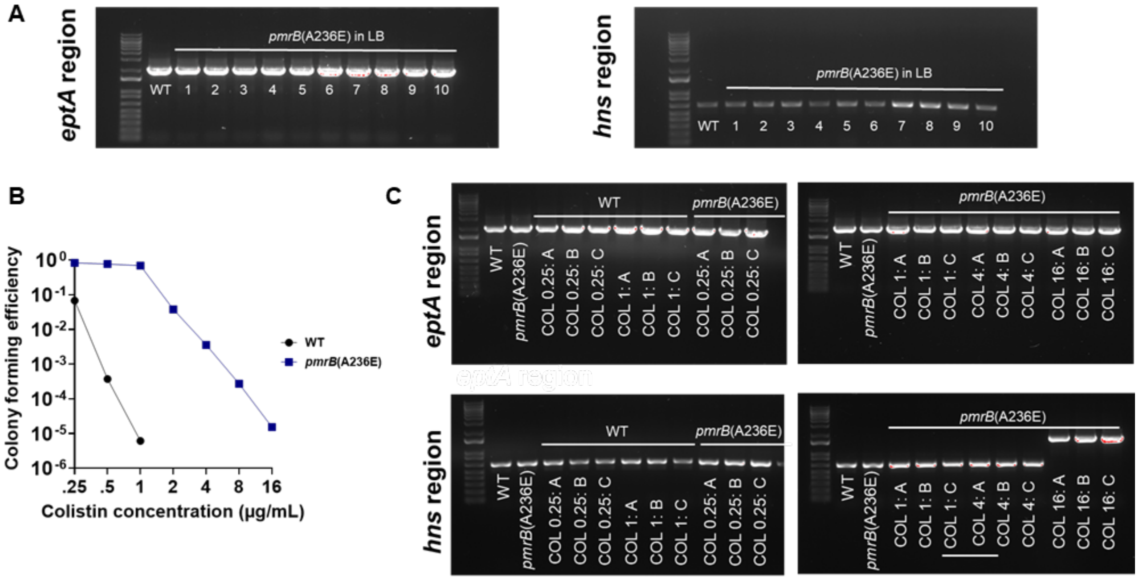


**Fig. S11. Insertion mutations in the vicinity of *eptA* or *hns* are only detected in the *pmrB*(A236E) mutant when plated at high colistin concentrations.**

**A.** The frozen stock of *pmrB*(A236E) was streaked on an LB plate. 10 individual colonies were randomly selected for colony PCR of the *eptA* and *hns* gene regions. WT was included as a negative (no insertion) control. **B.** Population analysis profiling of WT and *pmrB*(A236E)**.** Cultures of denoted strains are serially diluted and plated in LB supplemented with noted concentrations of colistin. Colony forming efficiency (CFE) was calculated as described. **C.** Colony PCR of the *eptA* and *hns* regions was performed on WT and *pmrB*(A236E) colonies selected at 0.25, 1, 4, and 16 µg/mL colistin (B). The presence or absence of insertion mutations was visualized by running the PCR products on a 1% agarose TAE gel. WT and untreated *pmrB*(A236E) were included as negative (no insertion) controls.

**Table S3. Strains used in this study.**

| Strain (clone name) | Description | Accession | COL MIC | Source | Relevant alleles |
|---|---|---|---|---|---|
| *E. coli* K12 | *E. coli* W3110 | - | - | (19) | - |
| ATCC17978 | *A. baumannii* ATCC17978UN | - | - | (20) | - |
| ATCC17978 R2 | Polymyxin-resistant ATCC17978UN | - | - | (20) | *pmrB*(T235I) |
| LAC-4 WT | Clinical isolate from Los Angeles County nosocomial outbreak. | SRX29091891, SRX29091953 | 0.38 | (21) | - |
| ΔpABLAC2 (Δ*GmR*) | Spontaneous plasmid loss resulting in Gentamycin sensitivity. | SRX29091892 | 0.38 | This | *ΔpABLAC2* |
| *pmrB*(T235I) (COL17) | COL 4 µg/mL selection from immune-depleted frozen pool (Line A, Passage 16) | SRX29091903, SRX29091954 | 16 | This | *pmrB*(T235I), ABLAC_16720::IS*Aba25*, ABLAC_23900::IS*Aba1*, ABLAC_35150::IS*Aba13* |
| *pmrB*(A236E) (COL23) | COL 2 µg/mL selection from Immunocompetent frozen pool (Line C, Passage 16) | SRX29091914, SRX29091955 | 0.75 | This | *pmrB*(A236E) |
| *lpxA** (LOS$^{(-)}$) | LOS deficient LAC-4. | SRX29091925 | 8 | This | *lpxA*(G142R) |
| *lpxA* pldA** (LB14) | LOS deficient LAC-4 selected at 42°C for improved growth. | SRX29091936 | 12 | This | *lpxA*(G142R), *ΔpABLAC2*, *pldA*(Δ1bp 84/1152). |
| *lpxA* pldA* ponA$^{ins}$* (LB13) | LOS deficient LAC-4 selected at 42°C for improved growth. | SRX29091947 | 12 | This | *lpxA*(G142R), *ΔpABLAC2*, *pldA*(Δ1bp 84/1152), *ponA*(+GAT 8/2466) |
| *lpxA* pldA* ponA** (LB14) | LOS deficient LAC-4 selected at 42°C for improved growth. | SRX29091958 | 12 | This | *lpxA*(G142R), *ΔpABLAC2*, *pldA*(Δ1bp 84/1152), *ponA*(E147Q) |
| A-0-Lo (I1) | Derived from COL23. COL 2µg/mL selection from Line A, Day 0. Colony collected from COL 16µg/mL plate. | SRX29091969, SRX29091956 | 6 | This | *pmrB*(A236E), new IS*Aba* |
| A-4-Lo (I2) | Derived from COL23. COL 2µg/mL selection from Line A, Day 4. Colony collected from COL 16µg/mL plate. | SRX29091977, SRX29091957 | 1 | This | *pmrB*(A236E), new IS*Aba* |
| A-7-Lo (I3) | Derived from COL23. COL 2µg/mL selection from Line A, Day 7. Colony collected from COL 16µg/mL plate. | SRX29091893, SRX29091959 | 2 | This | *pmrB*(A236E), *gacS*(310C), new IS*Aba* |
| B-0-Lo (I7) | Derived from COL23. COL 2µg/mL selection from Line B, Day 0. Colony collected from COL 16µg/mL plate. | SRX29091894, SRX29091960 | 2 | This | *pmrB*(A236E), new IS*Aba* |

| | | | | | |
|---|---|---|---|---|---|
| B-4-Lo (I8) | Derived from COL23. COL 2µg/mL selection from Line B, Day 4. Colony collected from COL 16µg/mL plate. | SRX29091895, SRX29091961 | 1.5 | This | *pmrB*(A236E), new IS*Aba* |
| B-7-Lo (I9) | Derived from COL23. COL 2µg/mL selection from Line B, Day 7. Colony collected from COL 16µg/mL plate. | SRX29091896, SRX29091962 | 6 | This | *pmrB*(A236E), *clpS*(Q90*), *eptA*(R127H) |
| C-0-Lo (I13) | Derived from COL23. COL 2µg/mL selection from Line C, Day 0. Colony collected from COL 16µg/mL plate. | SRX29091897, SRX29091963 | 3 | This | *pmrB*(A236E), new IS*Aba* |
| C-4-Lo (I14) | Derived from COL23. COL 2µg/mL selection from Line C, Day 4. Colony collected from COL 16µg/mL plate. | SRX29091898, SRX29091964 | 4 | This | *pmrB*(A236E), *clpA*(A302V), new IS*Aba* |
| C-7-Lo (I15) | Derived from COL23. COL 2µg/mL selection from Line C, Day 7. Colony collected from COL 16µg/mL plate. | SRX29091899, SRX29091965 | 4 | This | *pmrB*(A236E), new IS*Aba* |
| A-0-Hi (I4) | Derived from COL23. COL 4µg/mL selection from Line A, Day 0. Colony collected from COL 16µg/mL plate. | SRX29091900, SRX29091966, SRX29091976 | 6 | This | *pmrB*(A236E), new IS*Aba* |
| A-4-Hi (I5) | Derived from COL23. COL 4µg/mL selection from Line A, Day 4. Colony collected from COL 16µg/mL plate. | SRX29091901, SRX29091967 | 4 | This | *pmrB*(A236E), new IS*Aba* |
| A-7-Hi (I6) | Derived from COL23. COL 4µg/mL selection from Line A, Day 7. Colony collected from COL 16µg/mL plate. | SRX29091902, SRX29091968 | 4 | This | *pmrB*(A236E), new IS*Aba* |
| B-0-Hi (I10) | Derived from COL23. COL 4µg/mL selection from Line B, Day 0. Colony collected from COL 16µg/mL plate. | SRX29091904, SRX29091970 | 6 | This | *pmrB*(A236E), new IS*Aba* |
| B-4-Hi (I11) | Derived from COL23. COL 4µg/mL selection from Line B, Day 4. Colony collected from COL 16µg/mL plate. | SRX29091905, SRX29091971 | 6 | This | *pmrB*(A236E), *clpS*(+AC 132/414), new IS*Aba* |
| B-7-Hi (I12) | Derived from COL23. COL 4µg/mL selection from Line B, Day 7. Colony collected from COL 16µg/mL plate. | SRX29091906, SRX29091972 | 8 | This | *pmrB*(A236E), new IS*Aba* |

| | | | | | |
|---|---|---|---|---|---|
| C-0-Hi (I16) | Derived from COL23. COL 4µg/mL selection from Line C, Day 0. Colony collected from COL 16µg/mL plate. | SRX29091907, SRX29091973 | 1.5 | This | *pmrB*(A236E), new IS*Aba* |
| C-4-Hi (I17) | Derived from COL23. COL 4µg/mL selection from Line C, Day 4. Colony collected from COL 16µg/mL plate. | SRX29091908, SRX29091974 | 1.5 | This | *pmrB*(A236E), new IS*Aba* |
| C-7-Hi (I18) | Derived from COL23. COL 4µg/mL selection from Line C, Day 7. Colony collected from COL 16µg/mL plate. | SRX29091909, SRX29091975 | 2 | This | *pmrB*(A236E), ABLAC_02920(+T: -45nt upstream), new IS*Aba* |
| COL26 | COL 2 µg/mL selection from immunocompetent frozen pool (Line B, Passage 16) | SRX29091910 | 1.5 | This | *pmrA*(M12R) |
| COL27 | COL 4 µg/mL selection from immunocompetent frozen pool (Line B, Passage 16) | SRX29091911 | - | This | *pmrA*(M12R) |
| COL28 | COL 4 µg/mL selection from immunocompetent frozen pool (Line B, Passage 16) | SRX29091912 | 8 | This | *pmrB*(T235I), ABLAC_24300(P131S) |
| COL29 | COL 2 µg/mL selection from immune-depleted frozen pool (Line B, Passage 10) | SRX29091913 | 6 | This | *pmrB*(T235I), ABLAC_12660(Δ1bp 819/1569) |
| COL30 | COL 2 µg/mL selection from immune-depleted frozen pool (Line B, Passage 10) | SRX29091915 | 6 | This | *pmrB*(T235I) |
| COL31 | COL 4 µg/mL selection from immune-depleted frozen pool (Line B, Passage 10) | SRX29091916 | 6 | This | *pmrB*(T235I), ABLAC_06150(T15K) |
| COL33 | COL 1 µg/mL selection from immune-depleted frozen pool (Line C, Passage 10) | SRX29091917 | 0.19 | This | ISAba125 amplification, *envZ*(+GCCTTC 1084/1344), ABLAC_25350 (Δ1bp 290/387) |
| COL34 | COL 1 µg/mL selection from immune-depleted frozen pool (Line C, Passage 10) | SRX29091918 | 0.18 | This | *lpxC*(+GAATAT 501/903) |
| LB1 | LOS deficient LAC-4 selected at 42°C for improved growth. | SRX29091919 | - | This | *lpxA*(G142R), *ΔpABLAC2*, *pldA*(Δ1bp 84/1152), *ponA*(+GAT 8/2466) |
| LB2 | LOS deficient LAC-4 selected at 42°C for improved growth. | SRX29091920 | - | This | *lpxA*(G142R), *ΔpABLAC2*, *pldA*(Δ1bp 84/1152), *ponA*(E147Q) |
| LB3 | LOS deficient LAC-4 selected at 42°C for improved growth. | SRX29091921 | - | This | *lpxA*(G142R), *ΔpABLAC2*, *pldA*(Δ1bp 84/1152) |
| LB4 | LOS deficient LAC-4 selected at 42°C for improved growth. | SRX29091922 | - | This | *lpxA*(G142R), *ΔpABLAC2*, *pldA*(Δ1bp 84/1152), *ponA*(E147Q) |

| | | | | | |
|---|---|---|---|---|---|
| LB5 | LOS deficient LAC-4 selected at 42°C for improved growth. | - | - | This | *lpxA*(G142R) reverted to WT |
| LB6 | LOS deficient LAC-4 selected at 42°C for improved growth. | SRX29091923 | - | This | *lpxA*(G142R), *ΔpABLAC2*, *pldA*(Δ1bp 84/1152) |
| LB7 | LOS deficient LAC-4 selected at 42°C for improved growth. | SRX29091924 | - | This | *lpxA*(G142R), *msbA*(L161S), ABLAC_27930(K112*), ABLAC_09320( V54G), ABLAC_19710(K438N) |
| LB8 | LOS deficient LAC-4 selected at 42°C for improved growth. | SRX29091926 | - | This | *lpxA*(G142R), *ΔpABLAC2*, *pldA*(Δ1bp 84/1152) |
| LB9 | LOS deficient LAC-4 selected at 42°C for improved growth. | SRX29091927 | - | This | *lpxA*(G142R), *ΔpABLAC2*, *pldA*(Δ1bp 84/1152), ABLAC_25350(+TG 173/387) |
| LB10 | LOS deficient LAC-4 selected at 42°C for improved growth. | SRX29091928 | - | This | *lpxA*(G142R), *ΔpABLAC2*, *pldA*(Δ1bp 84/1152) |
| LB11 | LOS deficient LAC-4 selected at 42°C for improved growth. | SRX29091929 | - | This | *lpxA*(G142R), *ΔpABLAC2*, *pldA*(Δ1bp 84/1152), ABLAC_25350(+AA 55/387) |
| LB15 | LOS deficient LAC-4 selected at 42°C for improved growth. | SRX29091930 | - | This | *lpxA*(G142R), *ΔpABLAC2*, *pldA*(Δ1bp 84/1152), *ponA*(E147Q) |
| LB16 | LOS deficient LAC-4 selected at 42°C for improved growth. | - | - | This | *lpxA*(G142R) reverted to WT |
| LB17 | LOS deficient LAC-4 selected at 42°C for improved growth. | SRX29091931 | - | This | *lpxA*(G142R), *ΔpABLAC2*, *pldA*(Δ1bp 84/1152), *ponA*(E147Q) |
| LB18 | LOS deficient LAC-4 selected at 42°C for improved growth. | SRX29091932 | - | This | *lpxA*(G142R), *ΔpABLAC2*, *pldA*(Δ1bp 84/1152), *ponA*(E147Q) |
| PR1 (I4-1 S1) | A-0-Hi pseudorevertant selected on LB after drug free passaging (Day 14, Line 1) | SRX29091933, SRX30950811 | ColS | This | *pmrB*(A236E), new IS*Aba*, *pmrB*(Δ3bp: 1,206/1,335 nt) |
| PR2 (I4-1 S2) | A-0-Hi pseudorevertant selected on LB after drug free passaging (Day 14, Line 1) | SRX29091934 | ColS | This | *pmrB*(A236E), new IS*Aba*, *pmrB*(Δ3bp: 1,206/1,335 nt) |
| PR3 (I4-1 S3) | A-0-Hi pseudorevertant selected on LB after drug free passaging (Day 14, Line 1) | SRX29091935 | ColS | This | *pmrB*(A236E), new IS*Aba*, *pmrB*(Δ3bp: 1,206/1,335 nt) |
| PR4 (I4-1 S4) | A-0-Hi pseudorevertant selected on LB after drug free passaging (Day 14, Line 1) | SRX29091937 | ColS | This | *pmrB*(A236E), new IS*Aba*, *pmrB*(Δ3bp: 1,206/1,335 nt) |
| PR5 (I4-1 S5) | A-0-Hi pseudorevertant selected on LB after drug free passaging (Day 14, Line 1) | SRX29091938 | ColS | This | *pmrB*(A236E), new IS*Aba*, *pmrB*(Δ3bp: 1,206/1,335 nt) |

| | | | | | |
|---|---|---|---|---|---|
| PR6 (I4-1 S6) | A-0-Hi pseudorevertant selected on LB after drug free passaging (Day 14, Line 1) | SRX29091939 | ColS | This | *pmrB*(A236E), new IS*Aba*, *pmrB*(Δ3bp: 1,206/1,335 nt) |
| PR7 (I4-2 S1) | A-0-Hi pseudorevertant selected on LB after drug free passaging (Day 14, Line 2) | SRX29091940, SRX30950812 | ColS | This | *pmrB*(A236E), new IS*Aba*, *pmrB*(IS*Abe18*$^{ins}$: 163/1,335 nt) |
| PR8 (I4-2 S2) | A-0-Hi pseudorevertant selected on LB after drug free passaging (Day 14, Line 2) | SRX29091941, SRX30950814 | ColS | This | *pmrB*(A236E), new IS*Aba*, *pmrB*(G339V*) |
| PR9 (I4-2 S3) | A-0-Hi pseudorevertant selected on LB after drug free passaging (Day 14, Line 2) | SRX29091942 | ColS | This | *pmrB*(A236E), new IS*Aba*, *pmrB*(G339V*) |
| PR10 (I4-2 S4) | A-0-Hi pseudorevertant selected on LB after drug free passaging (Day 14, Line 2) | SRX29091943 | ColS | This | *pmrB*(A236E), new IS*Aba*, *pmrB*(G339V*) |
| PR11 (I4-2 S5) | A-0-Hi pseudorevertant selected on LB after drug free passaging (Day 14, Line 2) | SRX29091944 | ColS | This | *pmrB*(A236E), new IS*Aba*, *pmrB*(G339V*) |
| PR12 (I4-2 S6) | A-0-Hi pseudorevertant selected on LB after drug free passaging (Day 14, Line 2) | SRX29091945 | ColS | This | *pmrB*(A236E), new IS*Aba*, *pmrB*(G339V*) |
| PR13 (I4-3 S1) | A-0-Hi pseudorevertant selected on LB after drug free passaging (Day 14, Line 3) | SRX29091946, SRX30950815 | ColS | This | *pmrB*(A236E), new IS*Aba*, *pmrB*(Δ1bp: 1,299/1,335 nt) |
| PR14 (I4-3 S2) | A-0-Hi pseudorevertant selected on LB after drug free passaging (Day 14, Line 3) | SRX29091948, SRX30950816 | ColS | This | *pmrB*(A236E), new IS*Aba*, *pmrB*(Δ1bp: 1,299/1,335 nt) |
| PR15 (I4-3 S3) | A-0-Hi pseudorevertant selected on LB after drug free passaging (Day 14, Line 3) | SRX29091949, SRX30950817 | ColS | This | *pmrB*(A236E), new IS*Aba*, *pmrB*(Δ2bp: 1,121/1,335 nt) |
| PR16 (I4-3 S4) | A-0-Hi pseudorevertant selected on LB after drug free passaging (Day 14, Line 3) | SRX29091950 | ColS | This | *pmrB*(A236E), new IS*Aba*, *pmrB*(Δ1bp: 1,299/1,335 nt) |
| PR17 (I4-3 S5) | A-0-Hi pseudorevertant selected on LB after drug free passaging (Day 14, Line 3) | SRX29091951 | ColS | This | *pmrB*(A236E), new IS*Aba*, *pmrB*(Δ1bp: 1,299/1,335 nt) |
| PR18 (I4-3 S6) | A-0-Hi pseudorevertant selected on LB after drug free passaging (Day 14, Line 3) | SRX29091952, SRX30950818 | ColS | This | *pmrB*(A236E), new IS*Aba*, *pmrA*(IS*Aba25*$^{ins}$: 621/1,335 nt) |
| *pmrB*(T235I) BC | The *pmrB*(T235I) allele was backcrossed from COL17 into the WT strain. | SRX30950819, SRX30950821 | 4 | This | *pmrB*(T235I), new IS*Aba*, |
| *pmrB*(A236E) BC | The *pmrB*(A236E) allele was backcrossed from COL23 into the WT strain. | SRX30950820, SRX30950813 | 0.5 | This | *pmrB*(A236E) new IS*Abas* |

**Table S4. Primers and plasmids used in this study.**

| Primers | Sequence | Reference |
|---|---|---|
| **qRT_PCR primers** | | |
| 16s_qRT_For | CAGCTCGTGTCGTGAGATGT | (5) |
| 16s_qRT_Rev | CGTAAGGGCCATGATGACTT | (5) |
| pmrC_qRT_For | CCATTTGGCTAGGTGCAATTT | This |
| pmrC_qRT_Rev | ACCGCATAATAGGTAGCAACAA | This |
| naxD_qRT_For | ACCGATCAATACCCCTCATT | (22) |
| naxD_qRT_Rev | GTTCGCCATAATTGGTCAGT | (22) |
| eptA_qRT_For | TCTATCTGGCTAGGTTTATTTCTG | This |
| eptA_qRT_Rev | CTATTAAAATAACTAATGTCGCCCC | This |
| hns_qRT_For | CGGTAACGTGGTTCTACAGATTTA | This |
| hns_qRT_Rev | AAAGATCAAGCAATCGAAGATGC | This |
| weeH_qRT_For | GGATTACCCTGTGCATCAAATG | This |
| weeH_qRT_Rev | TGGGTAGCATATAAGGTTCGTAAA | This |
| ABLAC_36900_qRT_For | CCAAGGAGAGCAATAAGCAAAC | This |
| ABLAC_36900_qRT_Rev | GTGGTGGTGGAGTAGCAATAG | This |
| **Endpoint PCR** | | |
| hns_region_For | CGCTTGGCTTTTTAAGAGG | This |
| hns_region_Rev | TACAGGTGAAGCAGTTATGG | This |
| eptA_region_For | TCAACTGGGGATAACTATGC | This |
| eptA_region_Rev | GGCTCTAGATTACTCGAACC | This |
| pmrCAB_For | GTTTCAATTTCTACATTAAAGC | This |
| pmrCAB_Rev | TATTCATCGTTTTGAGTTC | This |
| ABLAC_35150_region_F | ATCGCTTGGCTTTTTAAGAG | This |
| ABLAC_35150_region_R | TACGACCAGAAAGTAGCATG | This |
| ABLAC_16720_region_F | GATGTATGAAGTCCGTCCAG | This |
| ABLAC_16720_region_R | AATATCTTTGGTAGCAGCCC | This |
| ABLAC_23900_region_F | TGAGTTTCTCTACACGCTTC | This |
| ABLAC_23900_region_R | CTTATGTTGTTACCTCCTGC | This |
| **Plasmid construction primers** | | |
| pmrB_Fw_NotI_800 | GACTCAGCGGCCGCAGTCAGGATAATTTATT | This |
| pmrB_BamHI_Rv_800 | CAGTCAGGATCCCTCGTCGCTTTTTTAT | This |

| Plasmids | Description | Reference |
|---|---|---|
| pJB4648APRA | derived from pSR47S. *oriTRP4 oriR6K sacB ApraR* | (4) |
| pJB4648APRA235-2-4 | pJB4648-NotI-*pmrBT235I*-BamHI | This |
| pJB4648APRA236-6-3 | pJB4648-NotI-*pmrBA236E*-BamHI | This |

### Dataset Legends

**Dataset S1 (separate file). Whole genome sequencing of mutants with altered colistin susceptibility in passaged pools.**

Isolates with altered colistin sensitivity able to grow on their respective colistin plates were saved and sequenced (Fig. S1). Colistin MIC of these isolates was determined via E-test strips (Materials and Methods). Mutations were detected by whole genome sequencing followed by BRESEQ analysis (Materials and Methods).

**Dataset S2 (separate file). Mutations present in LOS deficient suppressors selected at 42°C.**

LOS deficient isolates selected at 42°C in experiments described in Figure 3A were whole genome sequenced. Mutations were detected as described (Materials and Methods).

**Dataset S3 (separate file). Detection of IS*Aba* insertions in *pmrB*(A236E) isolates after selection for high colistin resistance.**

PacBio long-read sequencing was performed on denoted strains. Sequencing and alignment to LAC-4 chromosome performed as described (Materials and Methods). Identity of newly transposed IS*Aba* insertion sequences, their location and directionality are noted. The TA% of the genes and intergenic regions of insertion sites are noted.

**Dataset S4 (separate file). Mutations detected via the BRESEQ pipeline.**

Illumina short-read sequencing was performed on denoted strains. The BRESEQ pipeline was used to align reads to the reference genome and detect mutations. Only the mutations present in the evolved LAC-4 isolates and not their WT parent are shown. 100% means that all sequenced reads support the presence of the denoted mutation.
